# A neurogenetic toolkit to decode *Anopheles gambiae* olfaction

**DOI:** 10.1101/2023.08.16.553590

**Authors:** Diego Giraldo, Andrew M. Hammond, Jinling Wu, Brandon Feole, Noor Al-Saloum, Conor J. McMeniman

**Author notes:** These authors contributed equally (Diego Giraldo, Andrew M. Hammond).

## Abstract

The African malaria mosquito *Anopheles gambiae* exhibits a strong innate sensory drive to seek out human scent. To detect human odorants, *An. gambiae* uses olfactory sensory neurons (OSNs) that can be divided into different classes by unique repertoires of chemoreceptor gene expression. We applied CRISPR-Cas9-mediated T2A-In Frame Fusions and the *QF2/QUAS* system to gain genetic access to specific OSN subsets in *An. gambiae* expressing the chemoreceptor genes *Ir25a*, *Ir76b*, *Gr22* and *orco*. We first optimized methods to generate cell-type specific *QF2* driver and *QUAS* responder lines to map expression patterns of these chemoreceptors across mosquito sensory appendages. We next applied transcuticular calcium imaging to record neurophysiological responses to select human-related odorants for each OSN class. This neurogenetic toolkit tiling OSN subsets in *An. gambiae*, including those responsive to CO_2_, stands to support systematic efforts to decode olfaction in this prolific disease vector at high-resolution to combat malaria.

## Introduction

*Anopheles gambiae sensu stricto* (Diptera: Culicidae) is a devastating vector of human malaria throughout sub-Saharan Africa. This highly anthropophilic mosquito species blood feeds preferentially and frequently on humans (Besansky et al., 2004; Constantini et al., 1999; Dekker et al., 2001; Garrett-Jones et al., 1980; Scott and Takken, 2012; Takken and Verhulst, 2013) posing a major threat to public health in this region. To navigate towards a host from a range of distances *An. gambiae* females use their sense of smell to detect various chemical cues that signal human presence (Giraldo et al., 2023; Mukabana et al., 2002; Smallegange et al., 2010; Spitzen et al., 2013; Takken and Knols, 1999). Olfaction also plays a crucial role in other important *An. gambiae* behaviors such as nectar seeking (Foster and Takken, 2004; Nikbakhtzadeh et al., 2014; Nyasembe et al., 2012), detection of oviposition sites (Huang et al., 2006; Rinker et al., 2013), and swarm formation for mating (Mozūraitis et al., 2020).

The main olfactory appendages of female *An. gambiae* are the antennae, maxillary palps, and the labella of the proboscis. These organs house olfactory sensory neurons (OSNs) expressing chemosensory receptors such as odorant receptors (ORs), ionotropic receptors (IRs), and gustatory receptors (GRs) that bind volatile organic compounds encountered by *An. gambiae* in its sensory environment. Comparative genomics suggests that the *An. gambiae* genome putatively encodes 76 ORs, 110 IRs and 93 GRs (Matthews et al., 2018) – an unknown proportion of which are likely function in olfaction.

Studies of a subset of these ORs in *An. gambiae* and its closely related sibling species *An. coluzzii* have implicated this chemoreceptor gene family in detection of multiple classes of odorants such as aldehydes, ketones, esters, alcohols and aromatic compounds (Carey et al., 2010; Sun et al., 2020; Wang et al., 2010) that are present in volatile emissions from both host and nectar sources. Single ORs complex with the Odorant Receptor Co-Receptor (Orco) to form ligand-gated cation channels tuned to these ligands. In contrast, IRs in these two malaria vectors likely mediate sensation of amines and acids (Pitts et al., 2017; Raji et al., 2023) found predominantly in human scent; with each IR acting in conjunction with one or more IR co-receptors (IR8a, IR76b and IR25a) to detect odorants. A subset of GRs expressed in the *An. gambiae* and *An. coluzzii* maxillary palps (GR22, GR23 and GR24) have been found to form a CO_2_ receptor complex sensitive to this gaseous ligand (Lu et al., 2007, Liu et al., 2020; Omondi et al., 2015), that can also be activated by a range of other volatile organic compounds (Coutinho-Abreu et al., 2019).

Analyses in *Drosophila melanogaster*, *An. coluzzii* and *Aedes aegypti* have shown that while some OSNs only express one class of receptor, there are subsets of OSNs that express multiple, broadening their tuning properties (Herre et al., 2022; Raji et al., 2023; Task et al., 2022). This includes OSNs in these species that putatively co-express unique combinatorial repertoires of co-receptors such as Orco, IR76b, IR8a and IR25a alongside their associated ligand-tuning receptors. Relative to the above species, the molecular and cellular organization and odor-tuning dynamics of the *An. gambiae* olfactory system remains largely uncharacterized.

Recent advances in mosquito neurogenetics have been crucial to gain genetic access to subpopulations of OSNs using binary expression systems (e.g., QF2-QUAS system). This has facilitated neuroanatomical studies of the expression patterns of chemosensory receptors as well as their tuning properties (Afify et al., 2019; Herre et al., 2022; Raji et al., 2023; Ye et al., 2022; Zhao et al., 2021, 2022). Efforts to apply neurogenetic tools in *An. coluzzii* initially applied a promoter-fusion based strategy to characterize the expression pattern of the *orco* gene in the antennae, maxillary palps, and labella, and map the projections of *orco*+ OSNs into the antennal lobe (Riabinina et al., 2016). More recently in *An. coluzzii*, T2A-QF2 in-frame fusions have been applied to map the expression patterns of the *IR* co-receptor gene *Ir76b* in the antennae and labella (Ye et al., 2022), the tuning receptor gene *Ir41c* in multiple segments of the antenna (Raji et al., 2023), and the active zone protein gene *bruchpilot* throughout the nervous system (Konopka et al., 2023). Implementation of binary expression systems in this species has also enabled cell-type specific expression of genetically encoded calcium indicators (e.g., GCaMP6f) to study the tuning of Orco and IR41c expressing OSNs (Afify et al., 2019; Raji et al., 2023) by transcuticular calcium imaging. This approach enables rapid recording of odor-evoked responses from multiple individual OSNs at once in a non-invasive fashion, which makes it an ideal tool to screen for compounds that activate certain OSN classes in mosquitoes and study their ligand tuning dynamics.

Here, we engineer a neurogenetic toolkit to study the *An. gambiae* olfactory system. We report application of a two-step method to develop cell-type specific *T2A-QF2* driver lines for target *An. gambiae* chemoreceptors. Specifically, we demonstrate the utility of integrating a single multiplexed “driver-responder-marker” cassette consisting of a *T2A-QF2* driver directly coupled with a floxed *QUAS-mCD8::GFP* responder and an *Actin5C* (*Act5C*) fluorescent transgenesis marker to rapidly label OSNs expressing the olfactory chemoreceptor genes *Ir25a*, *Ir76b* and *Gr22* and *orco*. We highlight several factors for consideration when using such configurations to generate cell-type specific neurogenetic tools in *An. gambiae* including the potential for positional effects to modulate the expression of both responder and marker transgenes at these olfactory loci. We also demonstrate application of Cre-loxP excision to generate conventional *T2A-QF2* drivers for each target gene that are suitable for binary use, and map expression patterns across olfactory appendages using an *An. gambiae* responder line expressing the genetically encoded calcium indicator GCaMP6f (Chen et al., 2013). Finally, we apply transcuticular calcium imaging to record neurophysiological responses to select human-related odorants for each OSN class. These optimized transgenic lines tiling subsets of OSNs thus have applied utility for neuroanatomical and functional mapping of the *An. gambiae* olfactory system, and expand the neurogenetic toolkit available for molecular and cellular studies of olfaction in this primary vector of human malaria.

## Results

### A driver-responder-marker (DRM) system for rapid reporting of chemoreceptor gene expression patterns in the *Anopheles gambiae* olfactory system

T2A-In Frame Fusion technology has recently been deployed in both *An. gambiae* and *An. coluzzii* to map the expression patterns of a select handful of hygrosensors and thermoreceptors (Laursen et al., 2023), chemoreceptors (Raji et al., 2023; Ye et al., 2022) and pan-neuronal gene targets (Konopka et al., 2023). To date, all of these cell-type specific drivers have been developed by integrating a *T2A-QF2* transgene alongside an associated *3xP3* fluorescent transgenesis marker into the target locus via CRISPR-Cas9 mediated homology-directed repair (HDR). Given the need to select transformants, and then typically outcross and amplify the driver line before crossing an established responder line such as *QUAS-mCD8::GFP* from *An. coluzzii* (Riabinina et al., 2016) this process can typically take 3-4 generations before defining the neuroanatomical expression pattern of the target gene.

In this study, to rapidly report whether T2A-In Frame Fusions successfully gained genetic access to and labeled target subsets of OSNs in the *An. gambiae* olfactory system, we initially applied CRISPR-Cas9 mediated HDR to integrate a multiplexed transgenic cassette containing components of the *QF2/QUAS* binary expression system in-frame into coding exons of three olfactory co-receptor genes. These loci included *Ir76b* previously targeted to generate a complementary T2A-QF2 driver in *An. coluzzii* (Ye et al., 2022), as well as *Ir25a* and the CO_2_ receptor complex gene *Gr22* which have not previously been targeted to generate such reagents. In an attempt to develop lines that neuroanatomically report gene expression in a single transgenesis step visible in F_1_ transformants, we designed the integration cassette to consist of three parts: i) a *T2A-QF2* driver component, ii) a fluorescent *QUAS-mCD8::GFP* responder component that was floxed (flanked by *loxP* sites), and iii) an *Act5C*-based transgenesis marker component, forming a “driver-responder-marker” (DRM) system configuration (Figure 1A).

**Figure 1.**
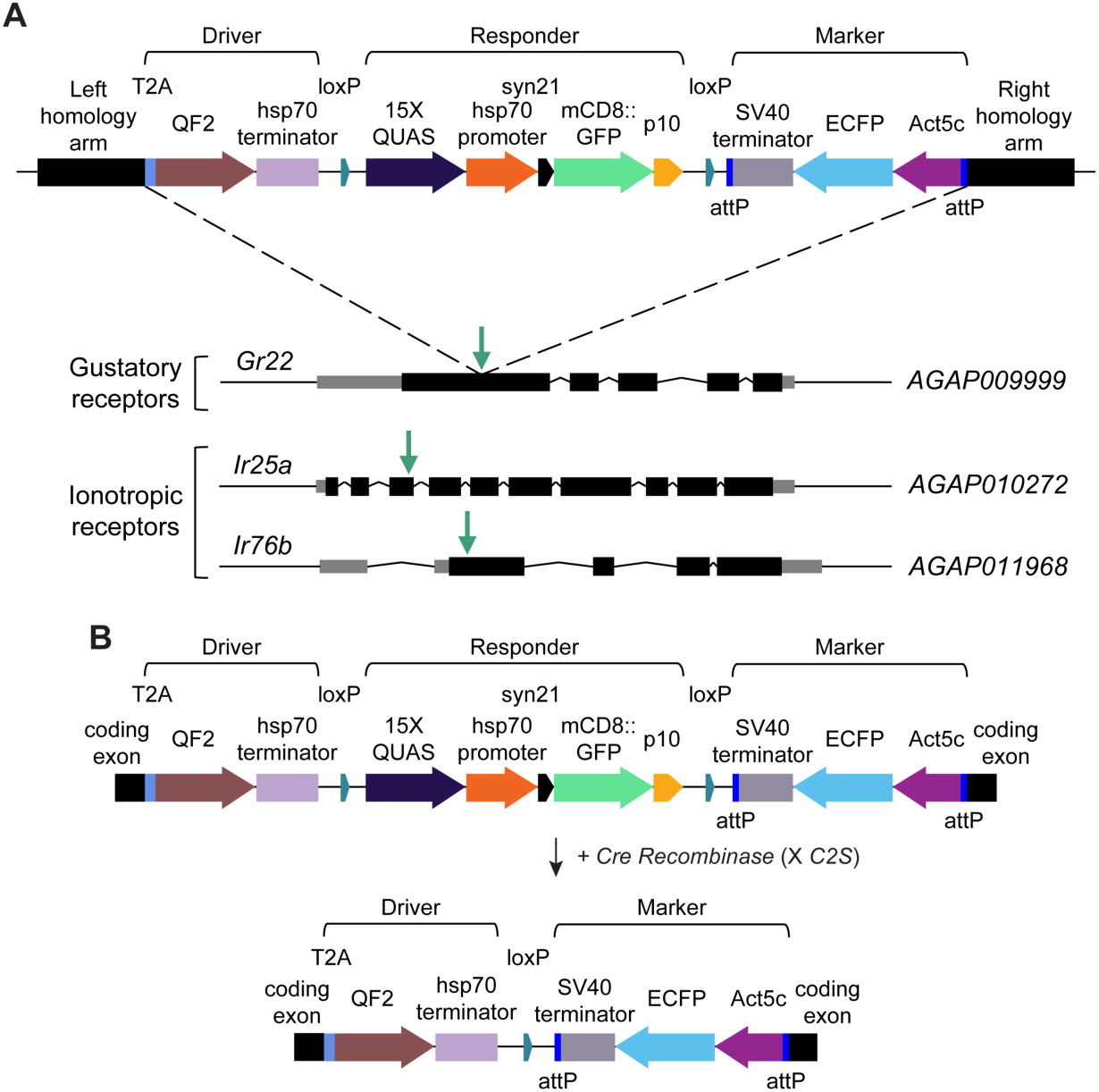
Schematic of the DRM system for rapid reporting of gene expression patterns and development of binary T2A-QF2 drivers for the *Anopheles gambiae* chemoreceptor genes *Gr22*, *Ir25a* and *Ir76b*. (A) Schematic of the DRM cassette integrated by CRISPR/Cas9-mediated HDR. The cassette contains three components: i) a *T2A-QF2* driver, ii) a *QUAS-mCD8::GFP* responder, and iii) an *Act5C-ECFP* transgenesis marker. These components are flanked by homology arms to catalyze in-frame insertion of this DRM cassette into coding exons of *Gr22*, *Ir25a* and *Ir76b* resulting in the lines *Gr22 ^DRM^, Ir76b ^DRM^*, and *Ir25a^DRM^*. (B) After DRM line establishment, the *QUAS-mCD8::GFP* responder component which is flanked by loxP sites can be removed via Cre-loxP mediated excision to generate conventional T2A-QF2 driver lines for binary use resulting in the lines *Gr22 ^QF2^, Ir25a^QF2^* and *Ir76b ^QF2^*. See also Figure S1.

Specifically, the “driver” consisted of a *T2A-QF2* transgene followed by a *Drosophila hsp70* terminator to drive the expression of the transcription factor QF2 under the control of the endogenous regulatory elements of each chemoreceptor gene when integrated in frame into a target coding exon. The “responder” consisted of a *15XQUAS-hsp70P-Syn21-mCD8::GFP-p10* transgene with a *Drosophila hsp70* core promoter (*hsp70P*) to drive the expression of mCD8::GFP expression in cell types where QF2 is expressed. We flanked this fluorescent reporter with both a *Syn21* sequence and *p10* 3’UTR shown to act as translational enhancers in *Drosophila* (Pfeiffer et al., 2012) to promote robust labeling of neuronal membrane and processes. We also strategically included *loxP*-sites surrounding this transgene to facilitate its prospective removal using Cre-loxP excision which is highly efficient in *An. gambiae* (Hoermann et al., 2021; Volohonsky et al., 2015) (Figure 1B). Finally, the “marker” consisted of an *Act5C-ECFP-SV40* transgene consisting of the *Drosophila Actin5C* (*Act5C*) promoter (Han et al., 1989) driving expression of enhanced cyan fluorescent protein (ECFP) with an SV40 terminator in the *An. gambiae* midgut to serve as a marker of successful transformation, and substitute for commonly used *3xP3* fluorescent markers which are also expressed in the nervous system (Laursen et al., 2023; Liu et al., 2020; Raji et al., 2023; Riabinina et al., 2016; Ye et al., 2022). The *Act5C-ECFP* marker was also surrounded by two φC31 attP sites to facilitate prospective exchange of this marker in lines post-establishment as desired using recombination-mediated cassette exchange (Bateman et al., 2006; Hammond et al., 2016; Adolfi et al., 2021).

For each target gene, we next designed a dual gRNA/Cas9 vector expressing a gRNA targeting a coding exon (Table S1), as well as a HDR donor plasmid containing the DRM cassette (Table S2) and microinjected these reagents into pre-blastoderm stage *An. gambiae* (G3 strain) embryos. Using this strategy, we successfully isolated F_1_ transformants for all three of these initial target genes by screening for visible expression of ECFP in the midgut of 1^st^ instar larvae. Subsequently, we outcrossed these lines to the wild type *G3* strain to generate the following DRM lines: *Gr22^DRM^*, *Ir25a^DRM^*and *Ir76b^DRM^*.

After line establishment, we then performed confocal analyses on the antennae, maxillary palps and labella (Figure 2A) of DRM females to determine the expression pattern of *Gr22, Ir76b*, and *Ir25a*. Strikingly, we observed expression of mCD8::GFP in all three appendages in these DRM lines (Figure S1). The expression was broad in *Ir25a^DRM^* with numerous cells strongly labeled with fluorescence in all three tissues (Figure S1A). *Ir76b^DRM^* also had broad expression in the labella and antennae, but only few fluorescent cells were found in the maxillary palps (Figure S1B). The *Gr22^DRM^* line showed expression in multiple cells of the maxillary palps, but only 1-2 cells in the antenna and labella (Figure S1C).

**Figure 2.**
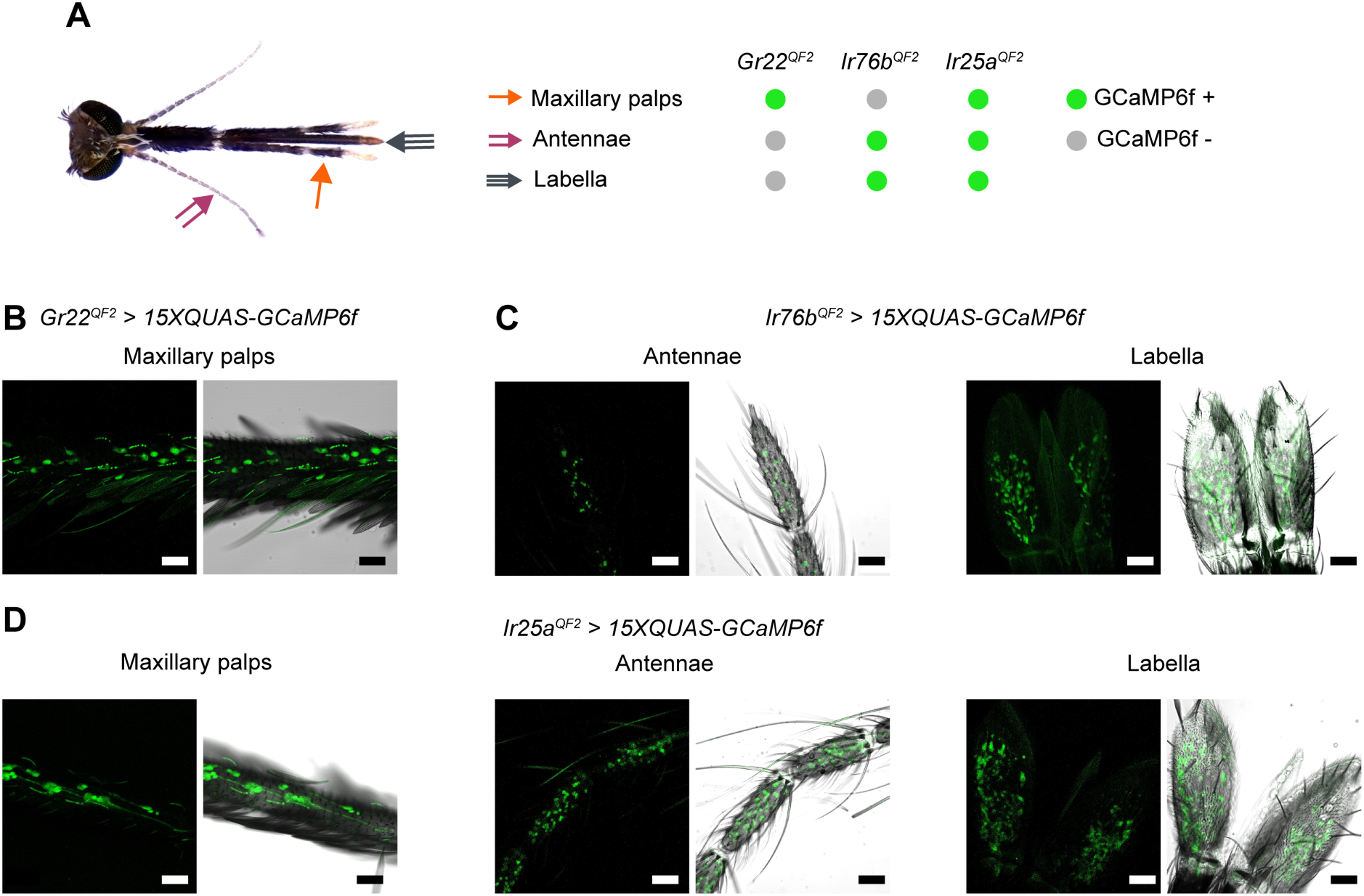
T2A-QF2 In Frame Fusions for binary use report expression patterns of the chemoreceptor genes *Gr22*, *Ir76b* and *Ir25a* in *Anopheles gambiae* olfactory appendages. (A) Image of the head of an *An. gambiae* female with the main olfactory appendages (antennae, maxillary palps, and labella of the proboscis) labeled. The right panel summarizes the observed pattern of *Gr22^QF2^*, *Ir76b^QF2^* and *Ir25a^QF2^*-transactivated *QUAS-GCaMP6f* expression in these appendages. (B) *Gr22*+ OSNs were found in the maxillary palps (genotype: *Gr22^QF2^, Act5C-ECFP > 15XQUAS-GCaMP6f, 3xP3-ECFP*). (C) *Ir76b*+ OSNs were restricted to the antennae and labella of the proboscis (genotype: *Ir76b^QF2^, Act5C-ECFP > 15XQUAS-GCaMP6f, 3xP3-ECFP*). (D) *Ir25a*+ OSNs were more broadly expressed in the maxillary palps, antennae, and labella (genotype: *Ir25a^QF2^, Act5C-ECFP > 15XQUAS-GCaMP6f, 3xP3-ECFP*). (B-D) Scale bar = 30µm. See also Figures 1, S2 and S3.

Transcriptomic analyses have revealed that *Gr22* is expressed by CO_2_-sensitive OSNs (cpA) found in the maxillary palps (Omondi et al., 2015) but not in the antenna (Rinker et al., 2013), which made expression of this gene in cells outside of the maxillary palps surprising. *Ir76b* transcripts have previously been detected in *An. coluzzii* maxillary palp tissue (Athrey et al., 2017; Pitts et al., 2011). However, a previous T2A-QF2 in-frame insertion in *Ir76b* in *An. coluzzii* reported expression of this gene in the antennae and labella, but not the maxillary palps (Ye et al., 2022).

To evaluate the potential for responder leakiness alone to generate such sparse labeling patterns in olfactory tissues, we next generated an independent *An. gambiae QUAS-mCD8::GFP* responder line using *piggyBac* mediated transposition (Figure S1D). This line included an *Act5C-DsRed* marker and gypsy insulators flanking the *QUAS-mCD8::GFP* cassette in an attempt to buffer any local cis-regulatory elements at the integration site, and enhancer elements associated with the *Act5C* promoter from influencing the expression of this responder transgene. Confocal analyses of olfactory appendages in a single isolated line carrying this responder transgene surprisingly revealed leaky mCD8::GFP expression in cells across the antennae, labella, and maxillary palps (Figure S1D) in the absence of QF2-based transactivation. This result suggests that some of the expression observed in these DRM lines may not be driven by the target genes – indicative of locus-dependent leaky expression (Figure S1B-D).

We conclude that the DRM system has the potential to rapidly report chemoreceptor gene expression patterns in the *An. gambiae* olfactory system. However, the potential for positional effects influencing the expression of responder transgenes within the context of the DRM system, as well as the potential for leaky expression in responder transgenes integrated elsewhere in the *An. gambiae* genome may confound interpretation of expression patterns in the absence of neurophysiological, transcriptomic or RNA in situ hybridization data for cross-validation.

### Cre-loxP excision of responder cassettes from *Anopheles gambiae* DRM lines generates conventional *T2A-QF2* drivers suitable for binary use in cell-type specific labeling

The DRM cassette we designed included a floxed *QUAS-mCD8::GFP* responder transgene (Figure 1A). To remove this fluorescent reporter we employed Cre-loxP cassette excision. To do this, we crossed each DRM line with the *C2S* line (Figure 1B) that expresses Cre recombinase in the *An. gambiae* germline (Hoermann et al., 2021; Volohonsky et al., 2015). We then selected individual progeny that did not have visible GFP labeling in adult olfactory tissue, but retained the *Act5C-ECFP* marker in the midgut; indicative of successful excision of the *QUAS-mCD8::GFP* responder. Confocal analyses of the antennae, labella, and maxillary palps from these responder excised lines showed no expression of mCD8::GFP in any cells across olfactory appendages (Figure S2A-C). We subsequently validated that these resulting lines only carried *T2A-QF2* driver and *Act5C-ECFP* marker transgenes with a single intervening loxP site using PCR (see Methods). We have denoted these conventional *T2A-QF2* driver lines as: *Gr22 ^QF2^, Ir76b* ^QF2^ and *Ir25a^QF2^*.

To determine the expression pattern of these target chemoreceptors in peripheral chemosensory appendages in the head of female *An. gambiae* we crossed *Gr22 ^QF2^, Ir76b* ^QF2^, and *Ir25a^QF2^* lines with an *An. gambiae QUAS-GCaMP6f* strain that we generated with *piggyBac*-mediated transposition using a construct previously validated in *An. coluzzii* (Afify et al., 2019). For these neuroanatomical studies, we used the fluorescent calcium indicator GCaMP6f (Chen et al., 2013) to label cells instead of our *QUAS-mCD8::GFP* line since we determined that the *An. gambiae QUAS-GCaMP6f* line that we generated lacked detectable leaky expression (Figure S1D and Figure S3). Confocal analyses revealed expression of GCaMP6f in maxillary palp OSNs of the *Gr22^QF2^ > QUAS-GCaMP6f* females (Figure 2B), as well as labeling of OSNs in the maxillary palps and labella of *Ir76b^QF2^ > QUAS-GCaMP6f* females (Figure 2C). We also found strong labeling in OSNs of the maxillary palps, antennae, and labella of *Ir25a^QF2^ > QUAS-GCaMP6f* females (Figure 2D). For each of these genotypes, we did not observe any fluorescence across any of the tissues sparsely labeled in the parental DRM integrations, or those carrying the *QF2* driver cassettes only (Figure S3). The loss of sparse labeling patterns in these binary *T2A-QF2* lines may thus result from excision of the *QUAS-mCD8::GFP* reporter, or alternatively the lower basal green fluorescence of GCaMP6f relative to mCD8::GFP making these rare cell types not visible.

### A DRM integration into the *Anopheles gambiae orco* gene illuminates positional effects influencing expression of both responder and marker transgenes

We hypothesized that the *Drosophila hsp70* core promoter used in both our initial DRM and independent *QUAS-mCD8::GFP* responder integrations may have been a potential source of reporter leakiness. To investigate alternative core promoters for use in the DRM system, we next replaced the *hsp70* core promoter associated with the *QUAS* elements of the DRM cassette (Figure 1A) with a *piggyBac* core promoter (Figure 3A) that has successfully been used for enhancer trapping in *An. stephensi* (O’Brochta et al., 2012) to generate a *15XQUAS-pBacP-Syn21-mCD8::GFP-p10* transgene. To pilot whether this transgene could label *An. gambiae* OSNs, we integrated this updated DRM system configuration in-frame into exon 2 of odorant receptor co-receptor *orco*, which is broadly expressed throughout the *An. gambiae* and *An. coluzzii* olfactory system (Pitts et al., 2011; Riabinina et al., 2016) using CRISPR-Cas9 mediated HDR.

**Figure 3.**
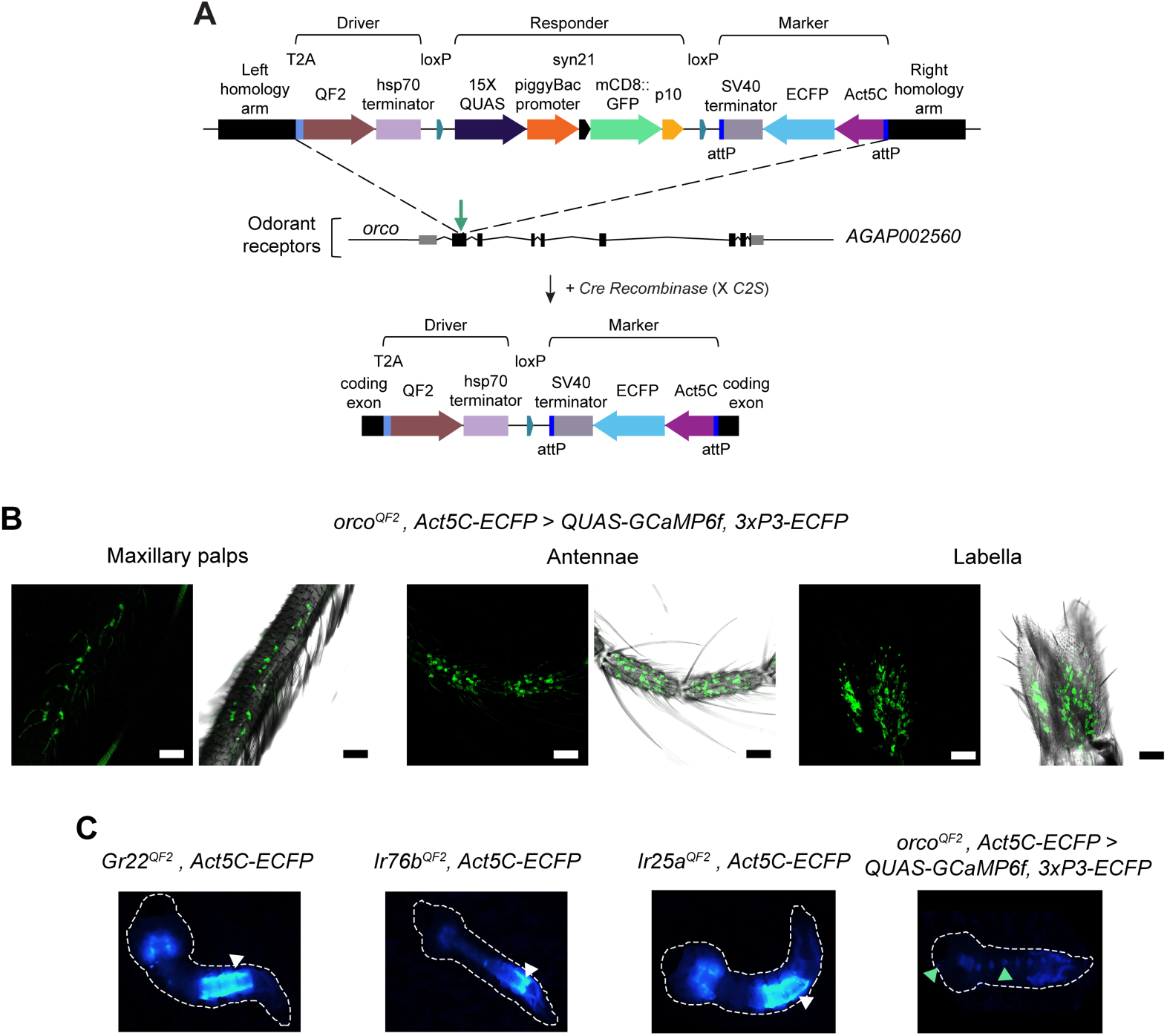
A DRM integration into the *Anopheles gambiae orco* gene reveals positional effects modulating responder and marker transgene expression. (A) Schematic of the DRM cassette for integration into the *An. gambiae orco* gene using CRISPR/Cas9-mediated HDR to generate *orco^DRM^*, and subsequent Cre-loxP excision of the responder transgene to generate *orco^QF2^* for binary use. (B) *orco*+ OSNs were found in the maxillary palp, antennae, and labella of the proboscis. (C) Locus-dependent heterogeneity in *Act5C-ECFP* marker expression across *An. gambiae* T2A-QF2 In Frame fusion lines. *Act5C-ECFP* expression (white arrows) in larval midgut marking *T2A-QF2* driver integration. *3xP3-ECFP* expression (green arrows) in ventral nerve cord and optic lobe marking *QUAS-GCaMP6f* responder integration. No visible expression of *Act5C-ECFP* was observed in the *orco^QF2^* integration, despite evidence of GFP labeling in OSNs in the larval antennae of both this cross and parental *orco^DRM^* line. See also Figures S4 and S5.

Notably, in F_1_ transformants we clearly observed mCD8::GFP expression in antennae of L1-L4 stage larvae (Figure S4A) indicating functionality of the *piggyBac* core promoter in this revised *QUAS-mCD8::GFP* responder transgene; facilitating the isolation of an *orco^DRM^*line. However, we did not observe any larvae expressing ECFP in the midgut, despite evidence of a fully intact *Act5C-ECFP* transgenesis marker being integrated in this line as revealed through Sanger sequencing (see Methods). Likewise, we did not observe any mCD8::GFP expression in adult chemosensory appendages suggesting the *piggyBac* core promoter may be active only in larval stages when integrated at this locus. To validate the integration of *T2A-QF2* into this location in *orco* was indeed capable of capturing endogenous regulatory elements suitable for reporting expression of this target gene across larval and adult stages of *An. gambiae*, we next excised this responder construct from the *orco^DRM^* by Cre-loxP excision as described above to generate a conventional *orco^QF2^* line (Figure 3A) for binary expression. We then crossed *orco^QF2^* with our *QUAS-GCaMP6f* line and in the resulting progeny found GCaMP6f expression in the larval antenna (Figure S4B) as well as strong labeling in OSNs in the antennae, labella, and maxillary palps (Figure 3B) of adult females.

These results indicate that the *piggyBac* core promoter is functional at *An. gambiae* larval stages for use in association with *QUAS* responder transgenes. However, we determined that this core promoter drives stage-stage specific responder transgene expression when integrated at the *orco* locus suggesting locus-dependent effects. Likewise, *Act5C-ECFP* when integrated in the *orco* locus also appears highly susceptible to positional effects as indicated by no visible expression of this marker in this genomic context. Nevertheless, larval specific mCD8::GFP expression within the context of a DRM configuration ensured successful isolation of initial transformants in the absence of a detectable *Act5C*-driven fluorescent transgenesis marker, facilitating the downstream establishment of a functional *orco^QF2^*line.

### Positional effects at *Anopheles gambiae Ir25a, Ir76b and Gr22* genomic loci influence the expression patterns of *Act5C* transgenesis markers

Consistent with positional effects influencing the expression pattern of the *Act5C-ECFP* marker when integrated into the *orco* gene, we noticed that the expression pattern of this fluorescent marker also varied across each of the other chemoreceptor genes that we targeted. For instance, in integrations generated at *Gr22* and *Ir25a* the *Act5C-ECFP* marker is expressed broadly throughout the midgut, whereas at *Ir76b* this marker is expressed in the posterior midgut (Figure 3C). Since *Act5C* is a commonly used promoter for *An. gambiae* and *An. coluzzii* transgenesis and considered a candidate substitute promoter to mark neurogenetic tools given its non-neuronal expression pattern, these positional effects influencing the activity of the *Act5C* promoter should be taken into consideration during construct design and screening to recover transformants for studies of *An. gambiae* neurobiology.

### T2A-QF2 In Frame Fusions facilitate transcuticular calcium imaging of OSN activity in *Anopheles gambiae* in response to stimulation with human-related odorants

Transcuticular calcium imaging has previously been applied to understand the ligand tuning dynamics of subsets of *An. coluzzii* OSNs expressing *orco* and *Ir41c* (Afify et al., 2019; Raji et al., 2023); and *An. gambiae* hygrosensitive neurons expressing *Ir93a* (Laursen et al., 2023). To validate the utility of the *T2A-QF2* lines developed here for this purpose, we next performed functional imaging with each of the *QF2* lines crossed to the *An. gambiae QUAS-GCaMP6f* line that we generated.

GR22 is a CO_2_ receptor complex subunit that is expressed in cpA neurons on the maxillary palps (Omondi et al., 2015) and this chemoreceptor is required for modulating sensitivity of this OSN class to CO_2_ and other odorants in *An. coluzzii* (Liu et al., 2020). We first tested whether *Gr22*+ OSNs found on the maxillary palp of *An. gambiae* females responded to CO_2_ by crossing our *Gr22^QF2^* driver with our *QUAS-GCaMP6f* responder, and imaging odor-evoked activity in maxillary palps of progeny carrying both of these transgenes (retaining one wild-type *Gr22* allele). We stimulated mosquitoes with a 1-second pulse of 1% CO_2_ and observed an increase in odor-evoked fluorescence in *Gr22*+ cells (Figure 3A). When we stimulated with increasing concentrations of CO_2,_ we found that odor-evoked responses increased with higher CO_2_ doses (Figure 3B-C). These results confirm that *Gr22*+ OSNs are indeed responsive to CO_2_.

To further validate transcuticular GCaMP imaging with *Ir76b^QF2^, Ir25a^QF2^ and orco^QF2^ > QUAS-GCaMP6f* females, we next stimulated antennal OSNs from each of these genotypes with ligands previously shown to activate these classes of chemoreceptors in *Drosophila* and *An. coluzzii* (Ni, 2021; Omondi et al., 2019), which are also found in human odor (Bernier et al., 2000; Martínez-Lozano, 2009; Rankin-Turner and McMeniman, 2022). We found that *Ir76b*+ OSNs in the 8^th^ antennal segment showed a significant response to a 0.28% trimethylamine stimulus relative to a water control (Wilcoxon signed rank test p = 0.0039, Figure 4A). Additionally, *Ir25a*+ OSNs in the 12^th^ antennal segment responded significantly more strongly to 1% pyridine relative to a water control (Wilcoxon signed rank test p = 0.002, Figure 4B) and *orco*+ cells in the same segment responded significantly more strongly to 1% sulcatone relative to a paraffin oil control (Wilcoxon signed rank test p = 0.002, Figure 4C).

**Figure 4.**
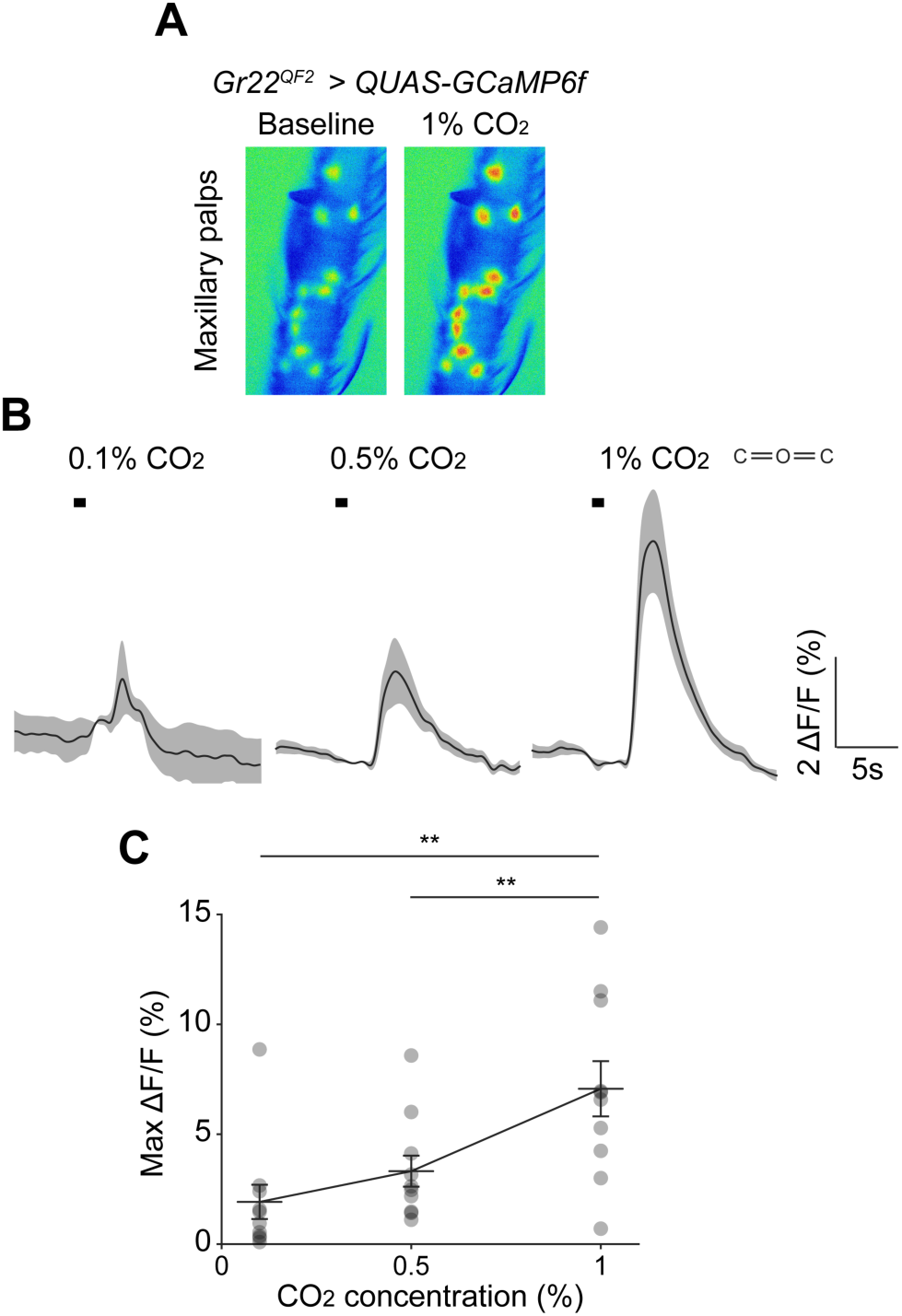
*Gr22*+ OSNs on the *Anopheles gambiae* maxillary palps respond to CO_2_. (A) GCaMP6f fluorescence in *Gr22* + OSNs before (left) and during (right) 1% CO_2_ stimulation (genotype: *Gr22^QF2^, Act5C-ECFP > 15XQUAS-GCaMP6f, 3xP3-ECFP*). (B) GCaMP6f traces of *Gr22*+ OSNs after 1s pulses (black bar) of increasing CO_2_ concentrations. Mean ±SEM plotted (n = 10). (C) Maximum change in fluorescence above baseline to increasing concentrations of 1s CO_2_ pulses. Mean ±SEM plotted (n = 10). Two-sided Wilcoxon signed rank test: 0% - 1% **p = 0.002; 0.5% - 1% **p = 0.0039.

**Figure 5.**
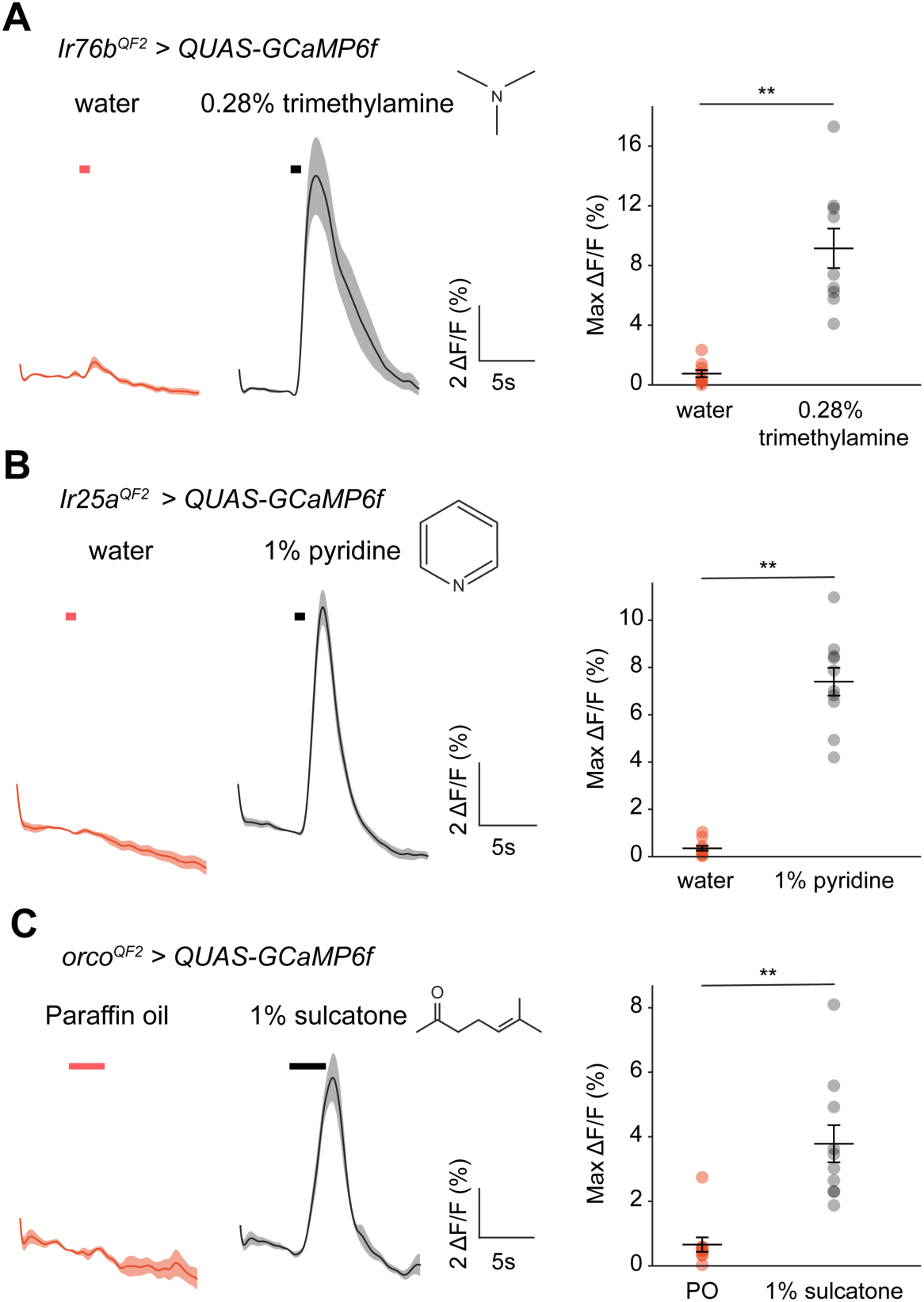
Odor-evoked activity of *Ir76b*+, *Ir25a*+ and *orco*+ OSNs in the *Anopheles gambiae* antennae to select human-related odorants. (A) GCaMP6f traces in *Ir76b*+ OSNs in the 8^th^ antennal segment in response to a 1s pulse of 0.28% trimethylamine or a water control (left) and quantification of the maximum ΔF/F values obtained from the traces (n = 9) for each stimulus (right). Genotype: *Ir76b^QF2^, Act5C-ECFP > 15XQUAS-GCaMP6f, 3xP3-ECFP*. (B) GCaMP6f traces in *Ir25a*+ OSNs in the 12^th^ antennal segment in response to a 1s pulse of 1% pyridine or a water control (left) and quantification of the maximum ΔF/F values (n = 10) (right). Genotype: *Ir25a^QF2^, Act5C-ECFP > 15XQUAS-GCaMP6f, 3xP3-ECFP*. (C) GCaMP6f traces of *orco*+ OSNs in the 12^th^ antennal segment in response to a 3s pulse of 1% sulcatone or a paraffin oil control (left) and quantification of the maximum ΔF/F values (n = 10) (right). Genotype: *orco^QF2^, Act5C-ECFP > 15XQUAS-GCaMP6f, 3xP3-ECFP*. (A-C) Orange and black bars indicate stimulus onset and duration. Mean ±SEM plotted in all panels. ** p < 0.01, two-sided Wilcoxon singed rank test.

We conclude that these T2A-QF2 lines validated here with transcuticular calcium imaging expand the repertoire of available tools in *An. gambiae* to study olfactory coding in OSN subsets expressing the chemoreceptor genes *Gr22*, *Ir76b*, *Ir25a* and *orco*.

## Discussion

Advances in our understanding of olfaction in malaria vectors have been spurred by integrative studies of their electrophysiological and behavioral responses to odorants (Braks et al., 2001; Cator et al., 2013; Cork and Park, 1996; Coutinho-Abreu et al., 2019; Knols et al., 1997; Lu et al., 2007; Meijerink and Van Loon, 1999; Omondi et al., 2015; Qiu et al., 2004, 2006; Takken et al., 2001), genomics and transcriptomics (Fox et al., 2001; Omondi et al., 2019; Pitts et al., 2011, 2017; Rinker et al., 2013), heterologous expression systems (Carey et al., 2010; Lu et al., 2007; Pitts et al., 2017; Wang et al., 2010) and genome editing (Liu et al., 2020; Sun et al., 2020). Most recently application of binary expression systems has provided insights into the neuroanatomy and function of OSNs and chemoreceptors, as well as other aspects of sensory biology of *An. gambiae* and *An. coluzzii.* These have included use of the QF2/QUAS system for cell-type specific labeling and calcium imaging using promoter fusion (Riabinina et al., 2016) and T2A-In Frame Fusion (Laursen et al., 2023; Raji et al., 2023; Ye et al., 2022, Konopka et al. 2023) -based QF2 drivers. To date, the vast majority of these neurogenetic tools targeting OSNs have been engineered in *An. coluzzii*. We therefore sought to expand the repertoire of neurogenetic tools available for studies of olfactory coding in *An. gambiae* given its status as a major primary vector of human malaria in sub-Saharan Africa.

Our approach to generate *QF2/QUAS* reagents in *An. gambiae* revealed several important considerations for engineering functional genetic tools in this non-model insect and related species. We initially developed a driver-responder-marker (DRM) system to rapidly obtain transgenic lines that report the expression pattern of chemoreceptor genes in the peripheral nervous system of *An. gambiae*, orthologous to recent designs in *Ae. aegypti* used to express genetically encoded calcium indicators pan-neuronally (Zhao et al., 2021) and in *orco*+ OSNs (Zhao et al., 2022). This system allowed us to, in a single step, insert a cassette that has a *T2A-QF2* driver, a floxed *QUAS-mCD8::GFP* responder and an *Act5C* fluorescent transgenesis marker using CRISPR-Cas9 mediated homologous recombination into target chemosensory genes. We initially piloted this approach using the co-receptor genes: *Gr22*, *Ir76b*, and *Ir25a*. We confirmed the DRM system provided a rapid method to label cells expressing these genes across olfactory appendages. These analyses revealed the *Ir25a^DRM^* lines broadly labeled cells in the antennae, maxillary palps and labella, *Gr22^DRM^*predominantly labeled cells in the maxillary palps and *Ir76b^DRM^* mostly labeled numerous cells in the antennae and labella; consistent with transcriptomic data indicating the expression of each of these genes in these olfactory tissues (Omondi et al., 2015; Rinker et al., 2013).

For the *Gr22^DRM^* line we also detected sparse cell labeling patterns in the antennae and labella that were discordant with current transcriptomic data indicating *Gr22* expression is restricted to the maxillary palps (Omondi et al., 2015; Rinker et al., 2013). For the *Ir76b^DRM^* line we conversely detected sparse labeling of cells in the maxillary palps which is supported by transcriptomic studies in *An. coluzzii* indicating the presence of *Ir76b* transcripts in this tissue (Athrey et al., 2017; Pitts et al., 2011). However, *Ir76b+* cells on the maxillary palps were not observed in a separate study using a complementary *Ir76b* T2A-QF2 *An. coluzzii* line and GFP labeling (Ye et al., 2022). Likewise, these sparse labeling patterns were not observed in the *Gr22^QF2^>QUAS-GCaMP6f* and *Ir76b^QF2^>QUAS-GCaMP6f* individuals we engineered and analyzed here. Whether such sparse labeling patterns found in our *Gr22^DRM^* and *Ir76b^DRM^* lines reflect leakiness of the *hsp70* core promoter associated with the *QUAS-mCD8::GFP* transgene in the DRM system configuration that we integrated into each of these loci, or these cells are labeled variably using these transgenic tools, or not always captured at the resolution of current transcriptomic analyses performed in these tissues awaits further characterization.

We also observed leaky reporter expression in numerous cells on the antenna, palp, and labella in the complete absence of QF2-based transactivation in an independent *An. gambiae QUAS-mCD8::GFP* responder line that we generated. We hypothesize that this undesirable labeling pattern may be observed in this line be due to local cis-regulatory elements acting on the *QUAS*-associated *hsp70* core promoter at the site of transgene integration yielding leaky reporter expression. We suggest such leaky reporter expression may also have been accentuated in this instance by the addition of gypsy insulators surrounding the responder cassette. Such an effect has been previously reported in *Drosophila* where insulator elements seem to have enhancer activity in certain tissues that lead to an increase in leaky expression of reporter genes under the control of UAS in a position dependent manner (Markstein et al., 2008). In an orthologous fashion, leakiness of reporter transgene expression in nervous tissues of a separate *QUAS-mCD8::GFP* responder line with a *hsp70* core promoter in *An. coluzzii* has been previously reported (Riabinina et al., 2016). These positional effects highlight the need to find loci in the *An. gambiae* genome where leakiness from QUAS or UAS responder transgenes is reduced or absent, as has been surveyed in *Drosophila* (Markstein et al., 2008; Pfeiffer et al., 2010).

We also determined using a DRM-based T2A-In Frame Fusion strategy in the *An. gambiae orco* gene that positional effects have the potential to influence the expression of both responder and marker transgenes. Specifically, we noted that a *piggyBac* core promoter we used in association with QUAS elements in the mCD8::GFP responder component of this line exhibited both an unexpected larval-specific labeling of OSNs that was absent in adults. We also found a complete absence of a visible *Act5C-ECFP* transgenesis marker in this line. The *piggyBac* core promoter has been previously used in Gal4-based enhancer trapping in adult *Anopheles stephensi* (O’Brochta et al., 2012) and therefore we anticipated it would be also active across life-cycle stages in *An. gambiae*. Likewise, we predicted the *Act5C-ECFP* marker would also be visible when integrated at this locus. Strikingly this was not the case in either instance, and future studies should therefore seek to identify basal core promoters that are not susceptible to such effects to ensure expression across life-cycle stages and to abrogate leakiness, as well as screen promoters for transgenesis markers that are best suited for each target locus. For instance, alternative synthetic core promoters such as the *Drosophila* synthetic core promoter have been used to replace the *hsp70* core promoter for binary expression systems in *Drosophila* (Pfeiffer et al., 2008, Wang et al., 2012), and endogenous core promoters have been employed with the GAL4/UAS system in *Tribolium castaneum* (Schinko et al., 2010). Likewise, systematic surveys in other *Drosophila* species have been used to identify new transgenesis markers suitable for use across different genomic loci (Stern, 2022).

While the DRM system is susceptible to positional effects, in the instance of the *orco* DRM integration it also proved to be an invaluable backup marker system where the *Act5-ECFP* maker used to report transgenesis was not active. The DRM approach is therefore highly useful when integrating cargo into genomic loci where the suitability of a chosen transgenesis marker for expression at that location is unknown, and the expression pattern of the target gene is predictable. For instance, in this study we tested the utility of the *Act5C* promoter to drive ECFP expression, which is canonically expressed in the *An. gambiae* midgut (Volohonsky et al., 2015), to confirm DRM integrations into target chemoreceptor genes. This was done in an attempt to investigate alternative promoters to replace the commonly used promoter *3xP3* (Berghammer et al., 1999) which can drive fluorescent marker expression variably throughout the nervous system (Stern, 2022). In the instance of the *orco* locus, we would have never recovered transformants without this DRM *QUAS-mCD8:GFP* responder transgene visibly labeling larval OSNs, due to the inactivity of the *Act5C* promoter at this genomic site.

To make DRM lines suitable for use in a binary fashion, we demonstrated that Cre-loxP excision (Volohonsky et al., 2015) can successfully convert these integrations into conventional *T2A-QF2* driver lines which have the potential to be crossed to any *QUAS*-responder transgene of interest. Using these reagents, when crossed with an *An. gambiae QUAS-GCaMP6f* line that we generated, we illuminated expression patterns and functional responses for each of our four target chemoreceptor genes. These results revealed that *Gr22^QF2^* drives GCaMP6f expression exclusively in OSNs of the maxillary palps where they respond to CO_2_ stimulation in a dose dependent manner. This was consistent with the known expression of *Gr22* in CO_2_ receptor complex neurons alongside *Gr23* and *Gr24* in *An. coluzzii* and *An. gambiae* maxillary palps (Lu et al., 2007; Liu et al., 2020). The *Ir76b^QF2^>QUAS-GCaMP6f* expression pattern we observed in the antennae and labella matched the transgenic labeling pattern previously reported for this gene in a *T2A-QF2* knockin in *An. coluzzii* (Ye et al., 2022). In this line we found OSNs in the antenna that responded to trimethylamine which is a ligand of OSNs expressing this coreceptor in flies (Vulpe & Menuz, 2021), and a known component of human odor (Martínez-Lozano, 2009).

In contrast, *Ir25a^QF2^>QUAS-GCaMP6f* mosquitoes exhibited a broader expression pattern across all olfactory appendages. Broad expression of *Ir25a* was also observed in *Ae. aegypti*, where it was found to be co-expressed with other receptor genes including *orco* and *Ir76b* (Herre et al., 2022). The broad expression pattern we observe for this receptor suggests that such co-expression in *An. gambiae* is highly likely. We specifically identified a subset of *Ir25a*+ antennal OSNs that responded to the aromatic compound pyridine found in human skin emanations (Bernier et al., 1999), which also is sensed by OSNs expressing this co-receptor in *Drosophila* (Ni, 2021). Finally, *orco^QF2^>QUAS-GCaMP6f* mosquitoes also revealed broad expression of this gene in all olfactory appendages, which is consistent with the expression pattern previously described using an *orco* promoter fusion *QF2* line of *An. coluzzii* (Riabinina et al., 2016). We identified *orco*+ OSNs on the antenna that responded to sulcatone, which is an abundant ketone present in human scent (Rankin-Turner and McMeniman, 2022).

These lines for transcuticular imaging therefore provide a rapid means to image odor-evoked activity in *An. gambiae*. These neurogenetic tools complement existing neurophysiological methods such as SSR (Coutinho-Abreu et al., 2019; Lu et al., 2007; Meijerink and Van Loon, 1999; Omondi et al., 2015; Qiu et al., 2006; Saveer et al., 2018) and EAG/EPG (Cator et al., 2013; Cork and Park, 1996; Knols et al., 1997; Qiu et al., 2004; Takken et al., 2001) to enhance studies of olfactory coding in these species. Future studies may therefore apply these reagents to comprehensively profile the ligand tuning dynamics of each of these OSN subtypes to additional components of human scent and other chemosensory stimuli.

In summary, we have successfully generated T2A-QF2 In Frame fusions in the chemoreceptor genes *Gr22*, *Ir25a*, *Ir76b* and *orco*, to provide genetic access to large subsets of OSNs collectively tiling the *An. gambiae* olfactory system. This work highlights several key considerations for engineering functional genetics tools in *An. gambiae* and related disease vectors, as well as non-model organisms. These validated reagents for neuroanatomy and functional imaging in *An. gambiae* expand the repertoire of neurogenetic tools available to study olfaction in this important malaria vector.

## Acknowledgements

This research was supported by funding from the Innovative Vector Control Consortium (P105) to C.J.M.; postdoctoral fellowships from the Human Frontier Science Program (LT000310/2019-L) and Johns Hopkins Malaria Research Institute to D.G.; and a Sir Henry Wellcome postdoctoral fellowship from the Wellcome Trust (213694/Z/18/Z) to A.M.H. Microscopy infrastructure at Johns Hopkins School of Medicine Microscope Core Facility used in this research was supported by the National Institutes of Health NCRR (S10OD023548). We thank Tony Southall, Stephen Goodwin and Andrea Crisanti for project advice and support; Kyros Kyrou, Eric Marois, Tony Nolan, Chris Potter and Gareth Lycett for constructs; Niyati Jain, James Tan, Matthew Gribble, Louise Marston, Ioanna Morianou, Ana Perez-Lebron, Margot Wohl and Stephanie Rankin-Turner for expert technical assistance; Olena Riabinina and members of the McMeniman lab for comments on the manuscript, and JHMRI and Bloomberg Philanthropies for generous supplemental funding and core infrastructure that supported this research.

## Author Contributions

Conceptualization, D.G., A.M.H. and C.J.M.; data curation, D.G. and J.W.; formal analysis, D.G., A.M.H., J.W. and C.J.M.; funding acquisition: D.G., A.M.H. and C.J.M.; investigation: D.G., A.M.H., J.W., B.F., N.A. and C.J.M.; methodology: D.G., A.M.H., J.W. and C.J.M.; project administration: C.J.M.; visualization: D.G. and J.W.; writing – original draft, D.G., A.M.H., J.W. and C.J.M.; writing – review & editing, all authors.

## Declaration of Interests

The authors declare no competing interests.

## Supplementary Figures

**Figure S1.**
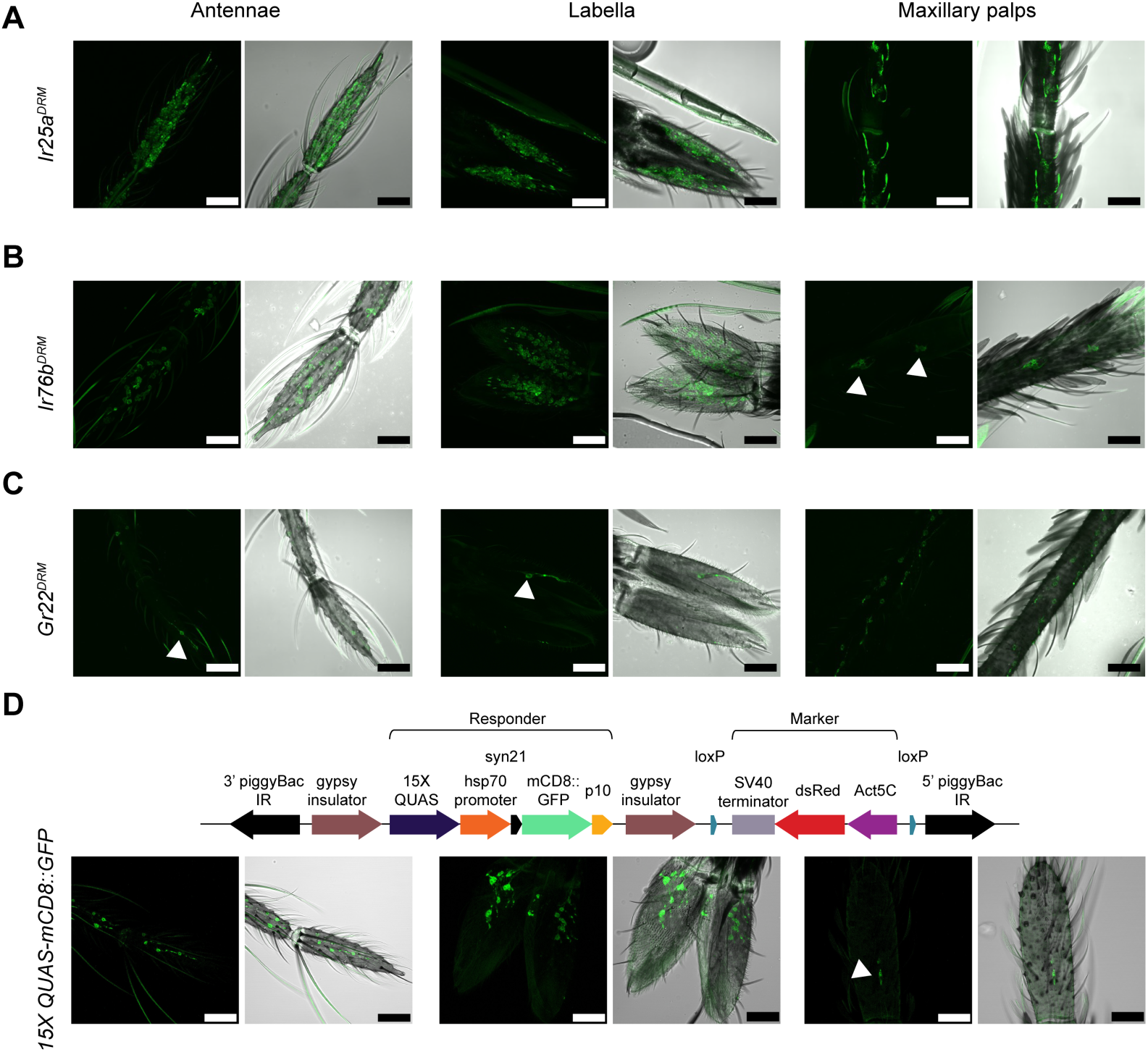
DRM integrations in *Ir25a*, *Ir76b* and *Gr22* rapidly report gene expression patterns across olfactory appendages, including sparse labeling of a few cells in some organs. Related to Figure 1. (A) *Ir25a^DRM^* broadly labels cells across the antennae, labellae and maxillary palps (genotype: *Ir25a^QF2^, QUAS-mCD8::GFP, Act5C-ECFP*). (B) *Ir76b^DRM^* broadly labels cells across the antennae and labella, and sparse cells in the maxillary palps (genotype: *Ir76b^QF2^, QUAS-mCD8::GFP, Act5C-ECFP*). (C) *Gr22^DRM^* broadly labels cells across the maxillary palps and sparse cells in the antennae and labella (genotype: *Gr22^QF2^, QUAS-mCD8::GFP, Act5C-ECFP*). (D) Schematic of QUAS-mCD8::GFP construct independently integrated into the *An. gambiae* genome using *piggyBac*-mediated transposition. The responder cassette was flanked by gypsy insulators. Leaky mCD8::GFP expression in the absence of QF2-based transactivation is observed in many cells in the antennae and labella and a few sparse cells in the maxillary palps. (E) (A-D) Scale bar = 30µm. (F) (B-D) Arrows indicate sparse labeling.

**Figure S2.**
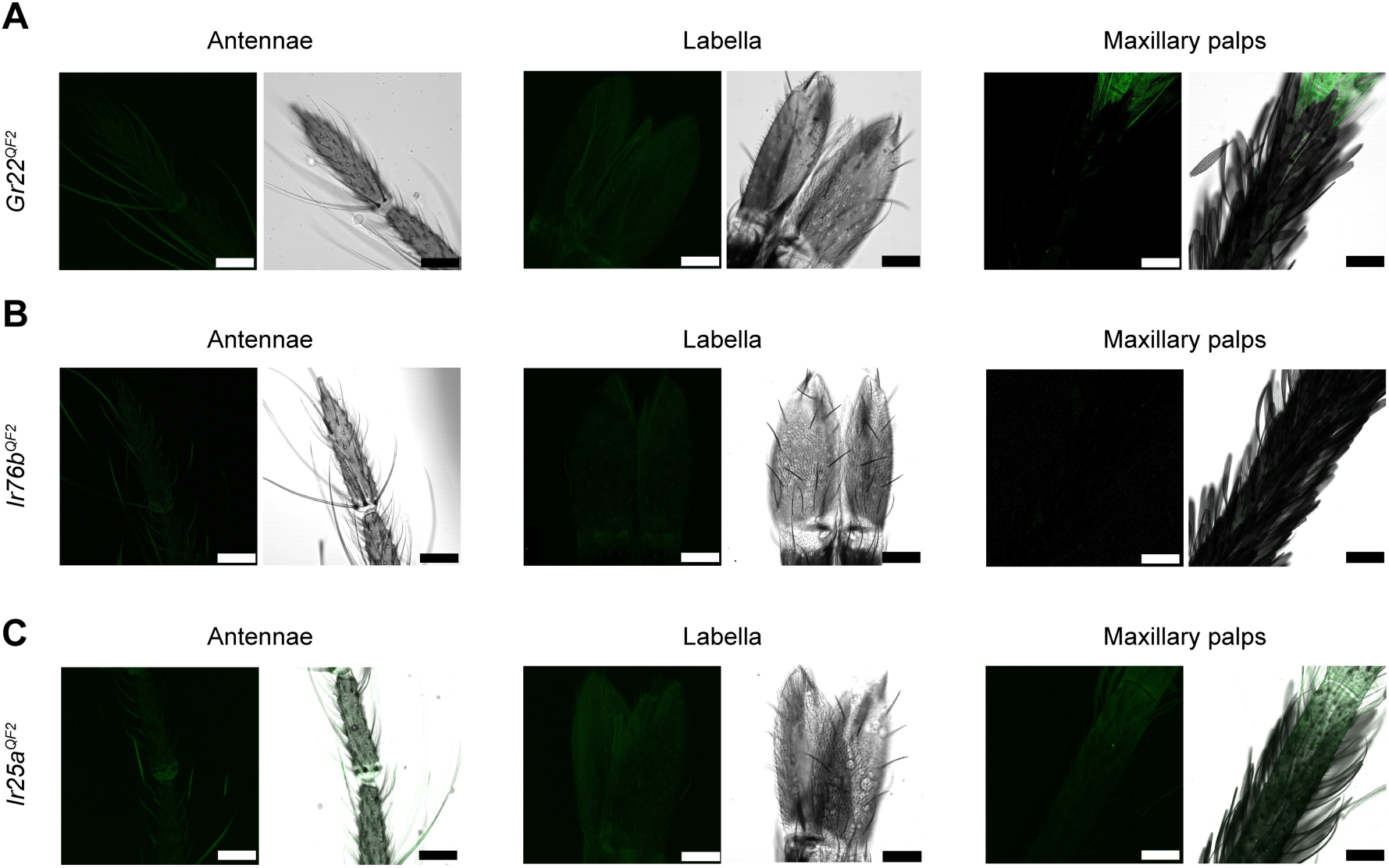
*Gr22*, *Ir76b* and *Ir25a* T2A-QF2 In Frame Fusions that are suitable for binary use show no evidence of background fluorescence in cells on *An. gambiae* olfactory appendages. Related to Figure 2. (A) *Gr22^QF2^* shows no evidence of cell labeling across the antennae, labella and maxillary palps (genotype: *Gr22^QF2^, Act5C-ECFP*). (B) *Ir76b^QF2^* shows no evidence of cell labeling across the antennae, labella and maxillary palps (genotype: *Ir76b^QF2^, Act5C-ECFP*). (C) *Ir25a^QF2^* shows no evidence of cell labeling across the antennae, labella and maxillary palps (genotype: *Ir25a^QF2^, Act5C-ECFP*). (A-C) Scale bar = 30µm.

**Figure S3.**
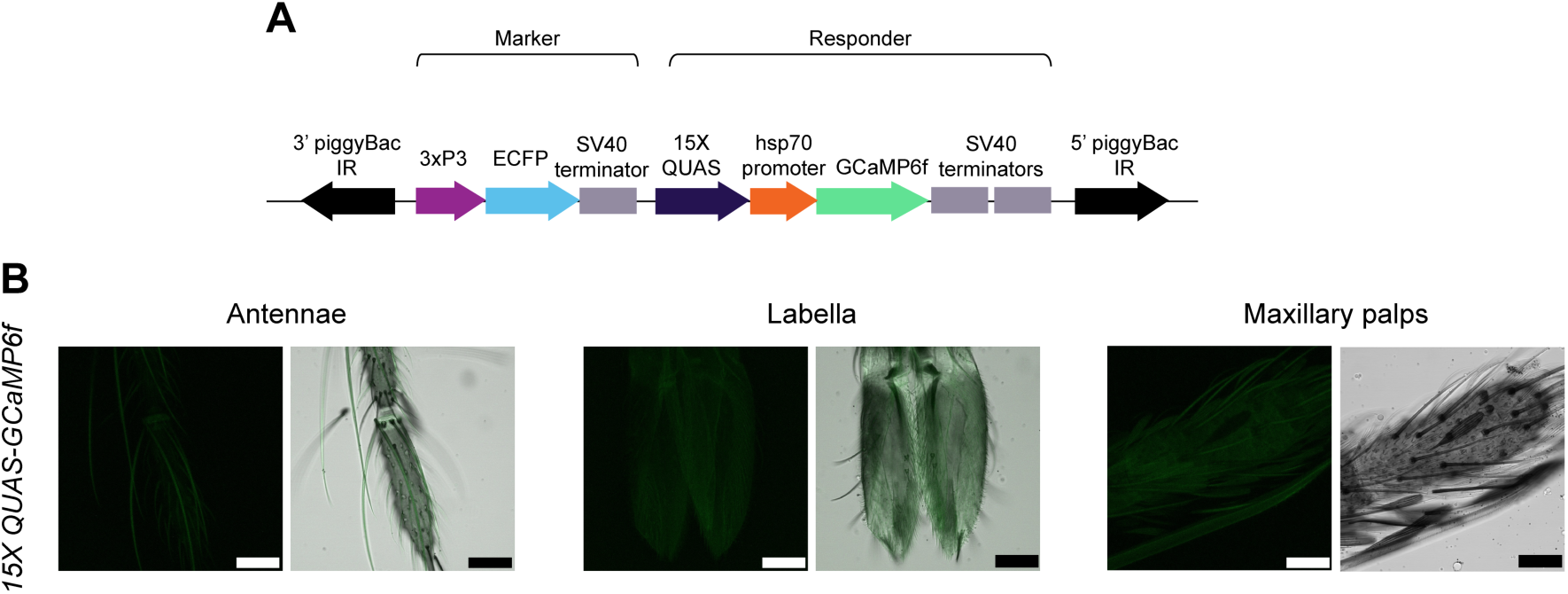
No evidence of visible background fluorescence in cells on the olfactory appendages of the *An. gambiae QUAS-GCaMP6f* line. Related to Figure 2. (A) Schematic of the *QUAS-GCaMP6f* construct used to generate an independent insertion of this transgene into the *An. gambiae* genome using *piggyBac* mediated transposition. (B) No GCaMP6f expression was visible across antennae, labella and maxillary palps of the *QUAS-GCaMP6f* responder line in the absence of *QF2*-based transactivation (genotype: *QUAS-GCaMP6f, 3xP3-ECFP*). Scale bar = 30µm.

**Figure S4.**
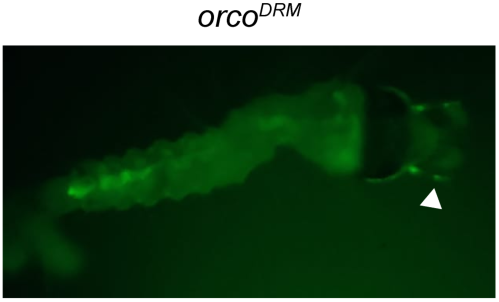
Antennal OSN labeling enables recovery of transformants during larval stages in a DRM integration into the *An. gambiae orco* gene. Related to Figure 3. A DRM integration into the *An. gambiae orco* gene (*orco^DRM^*) labeled antennal OSNs with mCD8::GFP that was visible from 1^st^ to 4^th^ larval instar stages. This *orco*+ labeling pattern facilitated the recovery of transgenics in the absence of a visible *Act5C-ECFP* expression marker in the larval midgut.

**Figure S5.**
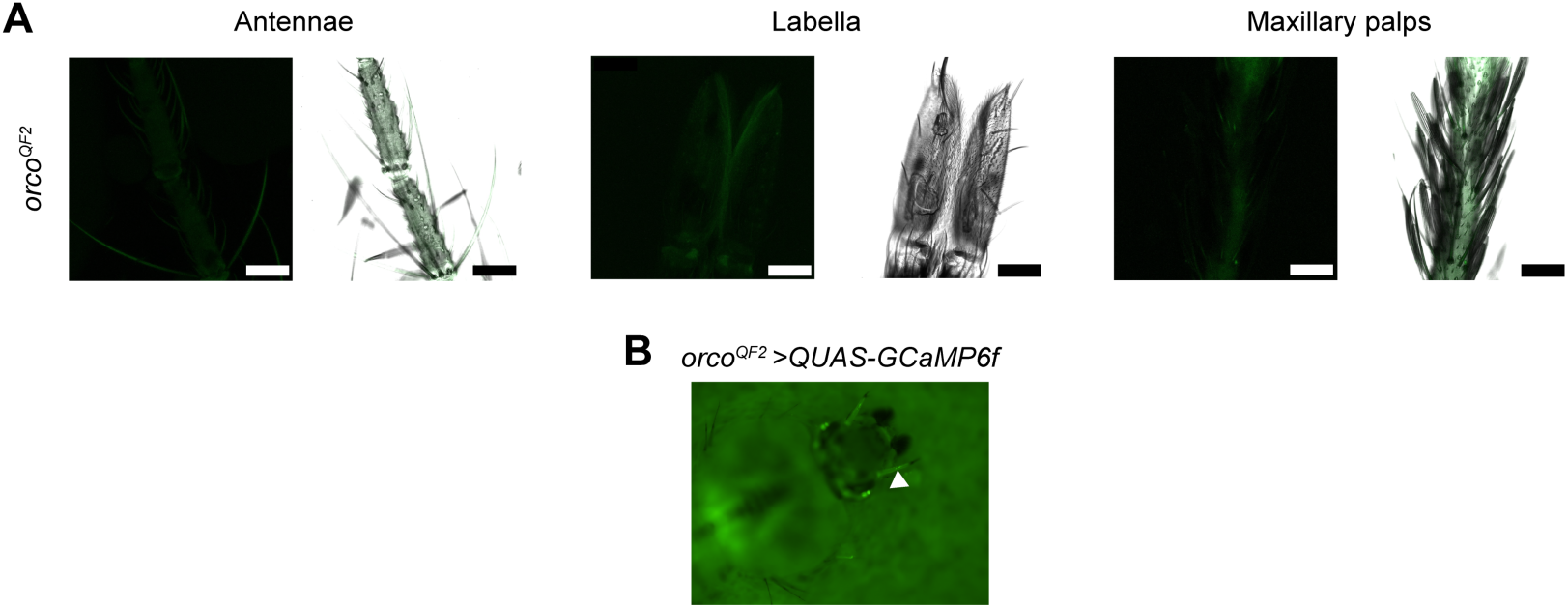
*orco* T2A-QF2 In Frame Fusion suitable for binary use shows no evidence of background fluorescence in cells on *An. gambiae* olfactory appendages and drives efficient expression of GCaMP6f in larval OSNs. Related to Figure 3. (A) *orco^QF2^* shows no evidence of background fluorescence in cells found across the antennae, labella and maxillary palps of adult females (genotype: *orco^QF2^, Act5C-ECFP*). (B) GCaMP6f expression is evident in *orco*+ OSNs of the larval antennae, and can be used to identify presence of the *orco^QF2^* driver in the absence of a visible *Act5C-ECFP* marker (genotype: *orco^QF2^, Act5C-ECFP > QUAS-GCaMP6f, 3xP3-ECFP*). Scale bar = 30µm. See also Figure 3C.

## Methods

### Mosquito strains and maintenance

The *Anopheles gambiae* G3, C2S (Volohonsky et al., 2015) and transgenic strains engineered in this study were maintained with a 14 hr light: 10 hr dark photoperiod at 27°C and 80% relative humidity using a standardized rearing protocol (Benedict et al., 2020). Adults were provided with a 10% sucrose solution for colony maintenance.

### gRNA/Cas9 expression constructs

The CRISPR vector p174 (Kyrou et al. 2018), expressing Cas9 under the *An. gambiae* promoter *zero population growth* and a gRNA under the U6 promoter, was secondarily modified by Golden Gate Cloning, as previously described (Kyrou et al. 2018) to contain gRNA spacers specific to targeting each of the olfactory target genes (Table S1). This yielded gRNA/Cas9 expression plasmids for *Ir25a* (p17424), *Ir76b* (p17422), *Gr22* (p17425) and *orco* (p17421).

### Generation of T2A-QF2 DRM constructs

A base HDR vector (p313) for integration of the driver-responder-marker (DRM) system into target *An. gambiae* chemoreceptor genes, was cloned using Gibson Assembly (New England Biolabs). Briefly, fragments containing a *T2A-QF2* driver (from pBB, Shankar et al., 2020), a *Syn21-15XQUAS-hsp70P-mCD8::GFP-p10* responder (from pMosECFP-15XQUAS-mCD8::GFP, Shankar et al., 2020) flanked by two loxP sites, and an *Act5C-ECFP* marker flanked by two φC31 attP sites (from pK104c, gift of K. Kyrou, Quinn et al., 2021) were assembled in the p174 vector (Kyrou et al., 2018) digested with AgeI and MluI. To target *Ir25a*, *Ir76b* and *Gr22*, p313 was secondarily modified to contain homology arms of approximately 1.5kb in length that were PCR amplified from *An. gambiae* G3 strain genomic DNA, extending outwards from each gRNA cleavage site (Table S2). The p313 base vector can be easily modified to contain any pair of homology arms in a single cloning step by digesting this vector with MluI and PmeI, co-purifying the two fragments in a single column cleanup step, and using Gibson Assembly to combine the two plasmid fragments with that of the homology arms at equimolar ratio. This assembly strategy yielded HDR vectors for *Ir25a* (p31304), *Ir76b* (p31302) and *Gr22* (p31305).

To target the *orco* gene, a modified DRM base plasmid (p327) was generated by substituting the *hsp70* core promoter in the responder component, with a *piggyBac* core promoter (*pBacP*) (O’Brochta et al., 2012) to generate a *15XQUAS-pBacP-Syn21-mCD8::GFP-p10* responder. Briefly, the fragment comprising *T2A-QF2-hsp70t-LoxP*, *15XQUAS*-*hsp70P* was excised from p313 by digestion with PmeI and XhoI. Gibson assembly of this digested p313 backbone, *T2A-QF2-hsp70t-LoxP* amplified from p313, and *QUAS-pBacP* sequence amplified from *pSL1180-QUAS-pBacP-mCD8GFP* (gift of D. Giraldo) was then performed. Homology arms for the *orco* target site were then amplified (Table S2) and inserted into p327 by digesting this base vector with MluI and PmeI and using Gibson assembly as described above for the other chemoreceptor genes, yielding the HDR vector for *orco* (p32701).

### Generation of independent responder constructs for insertion into the *An. gambiae* genome using piggyBac-mediated transposition

In Fusion cloning (Takara Bio) was used to assemble the responder vector (p371) for generation of an *An. gambiae QUAS-mCD8::GFP* line using *piggyBac*-mediated transposition. To generate p371, fragments comprising gypsy insulators from *Drosophila melanogaster*, *QUAS-hsp70P-syn21*, *HindIII-mCD8::GFP-p10*, and *LoxP-Act5C-DsRed* were PCR amplified (Table S3) and cloned into an empty p313 vector backbone, digested with PvuI and AscI. A derivative vector (p361) facilitates exchange of *mCD8::GFP* for alternative reporters by restriction digestion of this plasmid using HindIII and PmeI.

The plasmid pXL-BACII-ECFP-15xQUAS-TATA-Gcamp6f-SV40 (Afify et al, 2019) was used to generate an *An. gambiae QUAS-GCaMP6f* line in the G3 strain genetic background using *piggyBac*-mediated transposition.

### Microinjection

Pre-blastoderm stage embryos from the *An. gambiae* G3 strain were microinjected in the posterior pole as previously described (Fuch et al. 2013), using injection solutions comprising 300ng of each target-specific vector (i.e. HDR donor detailed in Table S1 and the corresponding Cas9/gRNA vector detailed in in 1x microinjection buffer (Fuch et al. 2013). Responder strains were created using injection solutions comprising 300 ng/µL each of the responder (i.e. p371 or pXL-BACII-ECFP-15xQUAS-TATA-Gcamp6f-SV40) and *vasa* transposase (pENTR R4-vas2-Transposase-R3) (Volohonsky et al., 2015) vectors. Briefly, freshly laid embryos were aligned against nitrocellulose paper and injected using a quartz capillary needle, three-axis micromanipulator (Narishige, MMO-4) and microinjector (Eppendorf FemtoJet 4X).

### Identification and verification of DRM integrations

*An. gambiae* G_0_ larvae with transient expression of fluorescent markers resulting from microinjection were isolated as virgin adults and outcrossed to the wild-type G3 strain. Transgenic F_1_ progeny with visible *Act5C-ECFP* expression in the larval midgut (*Ir25a^DRM^*, *Ir76b^DRM^* and *Gr22^DRM^*), or visible GFP expression in the larval antennae (*orco^DRM^*) were maintained as heterozygous lines by outcrossing as virgins to the G3 strain each generation. HDR cassette integration into each target gene was confirmed using PCR with a forward primer anchored outside of each left HDR arm and a reverse primer anchored in the QF2 coding sequence (Table S4).

### Cre-loxP excision of responder transgenes to yield T2A-QF2 In Frame fusions for binary use

To excise responder transgenes from each of the DRM strains, a minimum of 20 virgin females of the *C2S* line expressing Cre recombinase in the *An. gambiae* germline (Hoermann et al., 2021; Volohonsky et al., 2015) were mated to a minimum of 20 virgin males for each DRM line (i.e*. Ir25a^DRM^, IR76b^DRM^*, *Gr22^DRM^*and *orco^DRM^*). Progeny carrying both the *Act5C-ECFP* marker linked to the driver cassette, and *3xP3-DsRed* marker linked to the C2S line, were outcrossed to wild-type in reciprocal male and female crosses. Driver strains suitable for binary use (i.e. *Ir25a^QF2^, IR76b^QF2^*, *Gr22^QF2^* and *orco^QF2^*) were isolated by selecting progeny carrying the *Act5C-ECFP* marker linked to the driver cassette that also had no visible mCD8::GFP expression in larval OSNs (i.e. indicating successful excision of the floxed *QUAS-mCD8::GFP* responder); and which also lacked the *3xP3-DsRed* marker linked to *vas2-Cre* recombinase from the C2S line.

### Sequence verification of the *Act5C-ECFP* marker cassette in *orco^QF2^*

*orco^QF2^* driver mosquitoes were crossed with the *QUAS-GCaMP6f* responder line and adult F_1_ female progeny were screened for GCaMP6f fluorescence in the antenna and a 3xP3-ECFP maker to identify individuals carrying both of these transgenes (i.e. those with visible GFP labeling the larval antennae marking the presence of the *orco^QF2^* driver transgene and 3*xP3-ECFP* expression in larval optic nerves and ventral nerve cord marking the presence of the *QUAS-GCaMP6f* responder transgene). To confirm that the *Act5C-ECFP* marker associated with the *orco^QF2^* driver transgene (that was not visible) was fully intact, genomic DNA was extracted from a single F_1_ mosquito using the NucleoSpin Tissue kit (Macherey-Nagel Inc., 740952.50). A diagnostic PCR was then performed to fully amplify the *Act5C-ECFP* marker using the primer: 5’-TATCGATACCGTCGACTAAAGCC-3’ and 5’-CCAATTTGAAGTGCAGATAGCAGT-3’, and its sequence was verified using Sanger sequencing.

### Imaging of Peripheral Olfactory Appendages

Live antennal, palp and labella tissues were dissected in 0.1M PBS (119-069-131, Quality Biological) and immediately mounted in Slow-Fade Gold Antifade Mountant (Invitrogen, S36936) on glass slides. Images were acquired on a Zeiss LSM 880 Airyscan Fast confocal microscope within 2 hours of dissection. The 488 nm laser line was used to excite the green GCaMP6f signal. An additional DIC channel was used to visualize bright field morphology of the peripheral tissue. Images of the antennae, palps, and labella were acquired with a 40X oil immersion objective. Images were processed using Fiji (Schindelin et al., 2012). A maximum intensity projection was acquired for all z-slices of the GCaMP6f images.

### Mosquito preparation for peripheral calcium imaging

3-6 day old female mosquitoes were transferred to *Drosophila* vials (28.5 mm diameter, 95 mm long, 320110, Flystuff) containing a cotton ball soaked with dH_2_O for 1 hour before calcium imaging. Mosquitoes were cold anesthetized on ice for 3 minutes, and the wings and legs were gently removed. They were then placed sideways on double-sided tape (Scotch, 137DM-2) to affix the thorax and abdomen in position. The proboscis, antennae, and maxillary palps were placed on a coverslip to allow air to flow over them. These organs were pinned down with a quartz capillary (QF100-70-10, Sutter Instrument) to reduce movement of the preparation.

### CO_2_ delivery for calcium imaging

5% CO_2_ (X02AI95C2000117, Airgas) was mixed with clean air (UZ300, Airgas) using mass flow controllers (MC-series 100, MC-series 500, Alicat Scientific) to obtain the desired CO_2_ concentration at a flowrate of 2mL/s. The stimulus was pulsed by a solenoid valve (ETO-3-12, Clippard) controlled by an Arduino (Arduino UNO, Arduino) and a valve driver (ValveLink 8.2, Automate Scientific). These brought the CO_2_ stimulus into a carrier flow of humidified air (6mL/s) for a total flow of 8mL/s. A compensatory clean air flow (2mL/s) was on whenever the CO_2_ stimulus was off to ensure a constant flow of 8mL/s.

### Odorant delivery for calcium imaging

Trimethylamine (T0464, TCI) and pyridine (270407, Sigma-Aldrich) were diluted in molecular-grade dH_2_O to obtain a concentration of 0.28% and 1% respectively. Sulcatone (M48805, Sigma-Aldrich) was diluted in paraffin oil (PX0047-1, Supelco) to obtain a concentration of 1%. 100% dH_2_O or 100% paraffin oil were used as control stimuli. To prepare odor cartridges for stimulation, 40μL of solution were pipetted onto two round filter papers (Cytiva Whatman 2017006, 6 mm diameter, 20μL per paper) and the papers were inserted into 3cm segments of 1/4″ vinyl tubing (06405-02, Cole-Parmer). The cartridge was capped on both ends with parafilm until ready to use. Odors were delivered using a custom made olfactometer consisting of two manifold arrays (LFMX0510528B, The Lee Company) each containing 8 solenoid valves (LHDA1223411H, The Lee Company). Clean air (UZ300, Airgas) that was humidified was brought into a first manifold array at a flowrate of 2mL/s set with a mass flow controller (MC-series 100, Alicat Scientific). The odor cartridge was then connected to the output of one of the eight valves of the array on one end, and to the input of a solenoid valve in the second manifold array on the other end. The valves on each end of the odor cartridge were opened and closed simultaneously using an Arduino (Arduino UNO, Arduino) and a valve driver (ValveLink 8.2, Automate Scientific) to allow airflow over the headspace of the filter papers for odor stimulation. This system facilitates having up to 7 odor cartridges in the manifold array that can be independently controlled. The output of the second manifold array carrying the odor was connected to a tube that had carrier flow (6mL/s) of clean humidified air set by a mass flow controller (MC-series 500, Alicat Scientific) onto the live mosquito preparation. One of the channels of the manifold arrays was kept open between stimuli with humidified air flowing at 2mL/s and closed whenever the odor stimuli were on to ensure a constant flow of 8mL/s onto the preparation for the duration of the experiment. The odor cartridges were replaced between each mosquito preparation.

### Calcium imaging system and analysis

Calcium signals were recorded using an epifluorescence microscope (BX51W1, Olympus) and an LED light source (Lambda HPX-L5, Sutter Instrument) and a 50X objective (LMPlanFLN, Olympus, 0.5 NA). Recordings were carried out at 30 fps using a CMOS camera (Orca-Fusion C14440, Hamamatsu Photonics K.K.) controlled by Micro-Manager (Edelstein et al., 2014). The videos were analyzed using FIJI (Schindelin et al., 2012) to obtain ΔF/F_o_ traces. Regions of interest (ROIs) were selected using the “ROI manager” tool in FIJI, and fluorescence intensity traces were obtained with the multi measure function. For videos with CO_2_ stimulation of *Gr22^QF2^>QUAS-GCaMP6f* mosquitoes, an ROI was drawn around every fluorescent cell body in the fourth segment of the palp, and the responses of all units were averaged for each mosquito preparation. For videos of odor-evoked responses from *Ir76b^QF2^>QUAS-GCaMP6f*, *Ir25a^QF2^>QUAS-GCaMP6f* and *orco^QF2^>QUAS-GCaMP6f* mosquitoes, ROIs were drawn around cell bodies that were visibly responding to each stimulus in the specified antennal segment, and the response of these units was averaged for each preparation. The analysis of the calcium traces obtained from FIJI and all statistical analyses were performed using Matlab R2018b (The MathWorks Inc.). To calculate ΔF/F_o_, the F_o_ value was calculated by averaging the fluorescence intensity of all the frames 0.5 seconds before stimulus onset.

## QUANTIFICATION AND STATISTICAL ANALYSIS

We tested normality of maximum responses to treatment and control stimuli in calcium imaging experiments with a Kolmogorov-Smirnov test. To test for the statistical significance of differences observed we used a Wilcoxon signed-rank test. All statistical analyses we performed in Matlab R2018b (The MathWorks Inc.).

**Table S1.**
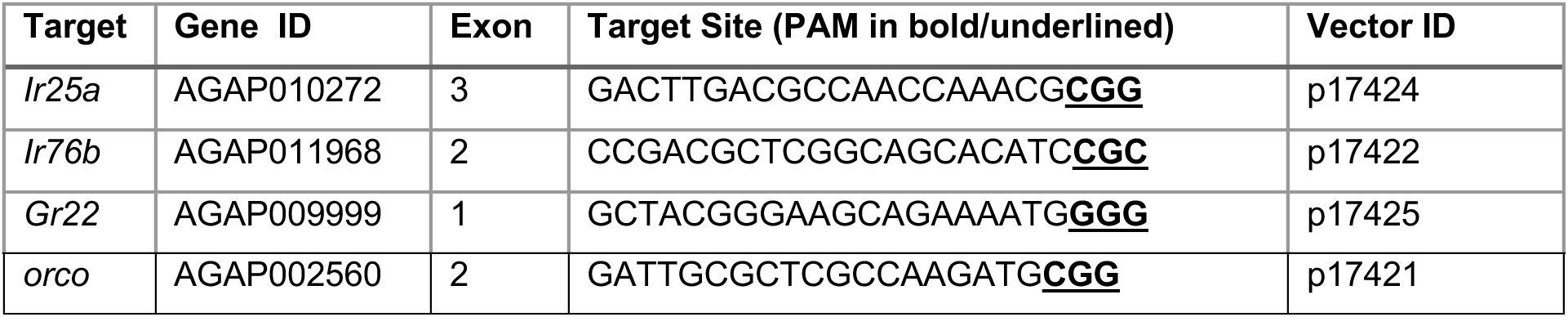
Sequences of gRNA target sites for *Anopheles gambiae* chemoreceptor genes and the resulting vector IDs used for gRNA/Cas9 expression. Related to Methods.

**Table S2.**
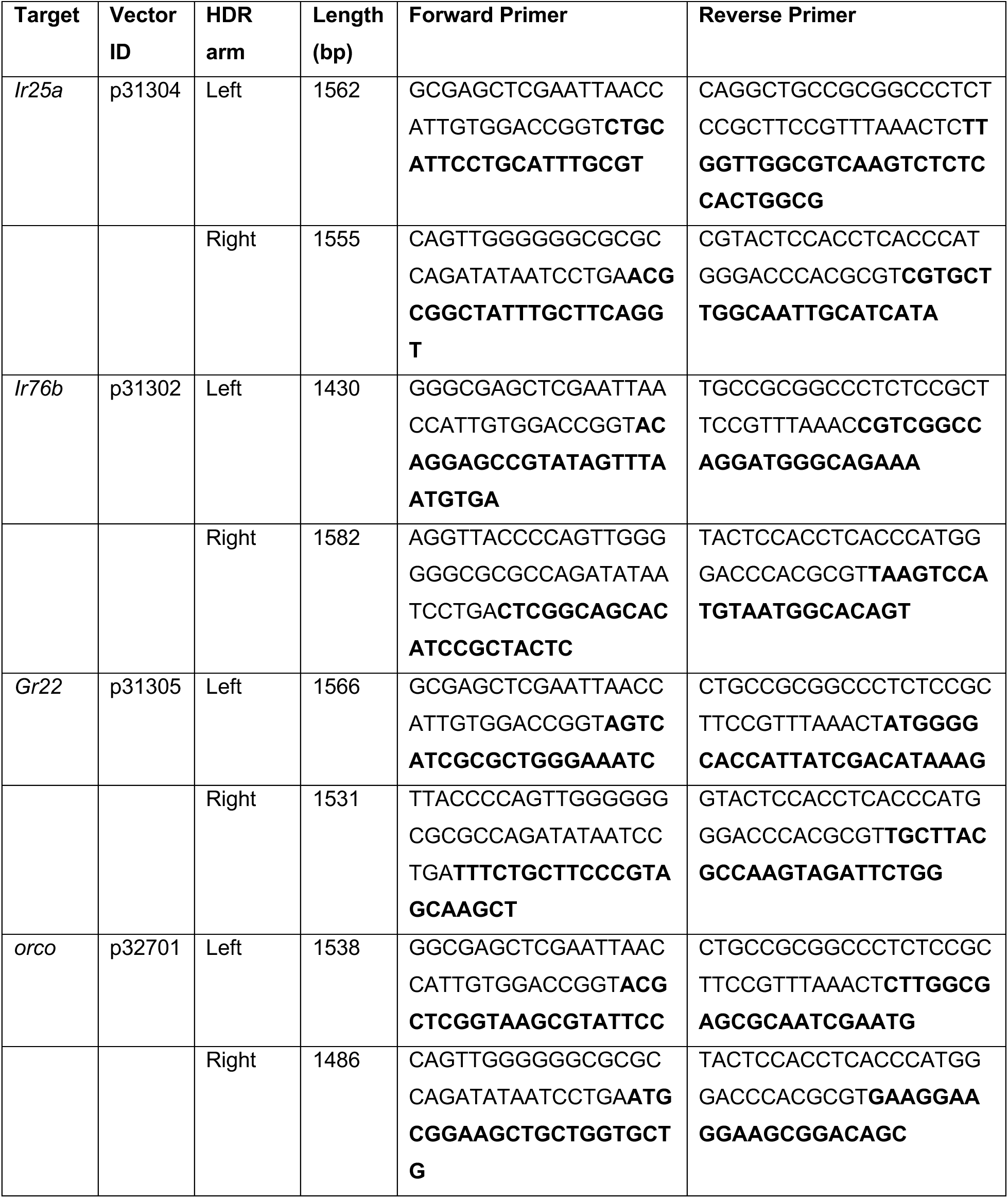
HDR vector IDs and primers used to amplify homology arms for DRM system integration into *Anopheles gambiae* chemoreceptor genes. Primers comprise genome-specific sequences (in bold) and 5’ adaptors to mediate insertion by Gibson Assembly into p313 (*Ir25a*, *Ir76b* and *Gr22*) or p327 (*orco*) base vectors. Related to Methods.

**Table S3.**
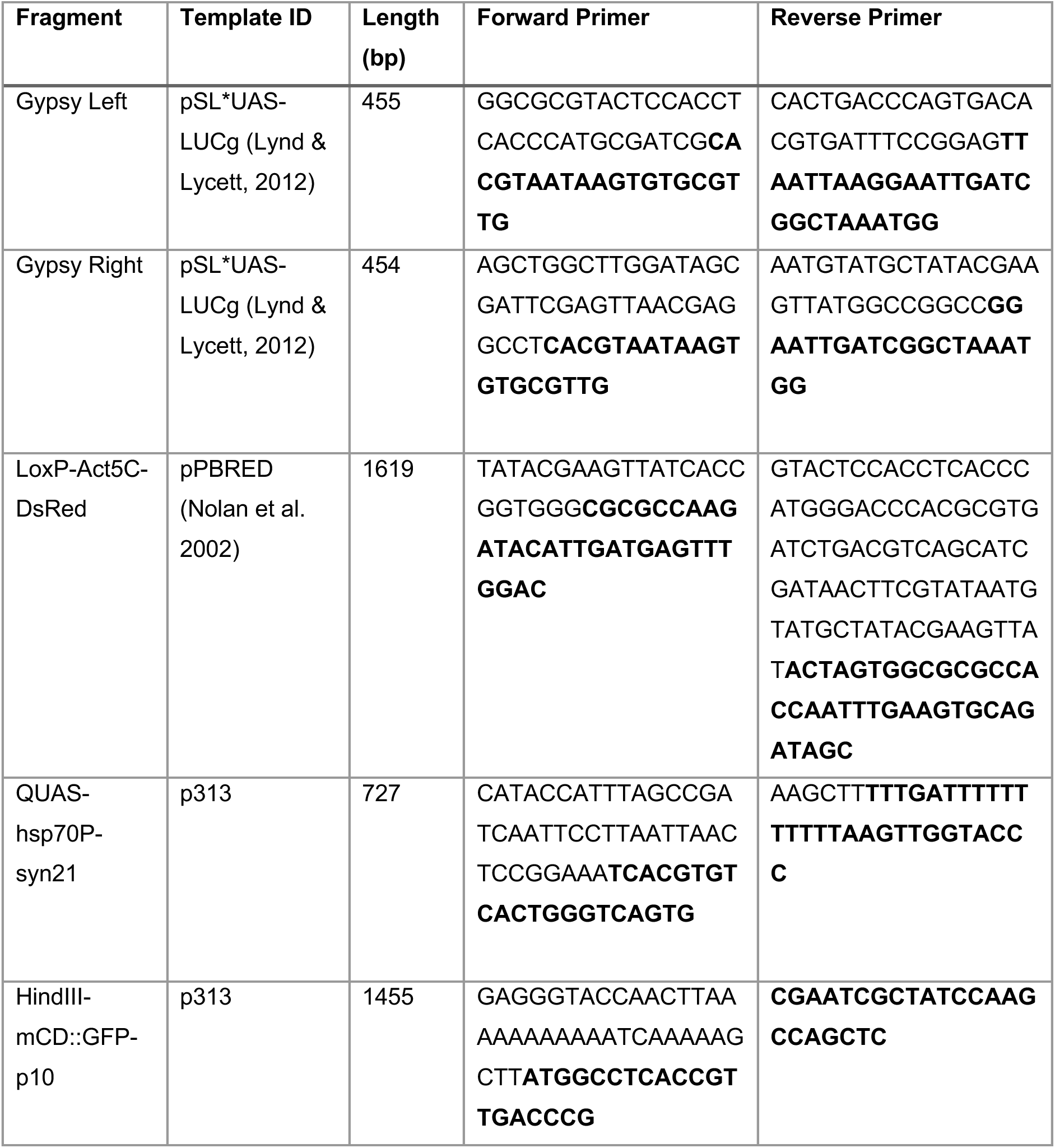
PCR primers and templates used for In Fusion cloning to assemble the vector p371 used to generate the *Anopheles gambiae QUAS-mCD8::GFP* line via *piggyBac*-mediated transposition. Template specific primers are highlighted in bold, with 5’ InFusion adapters. Related to Methods.

**Table S4.**
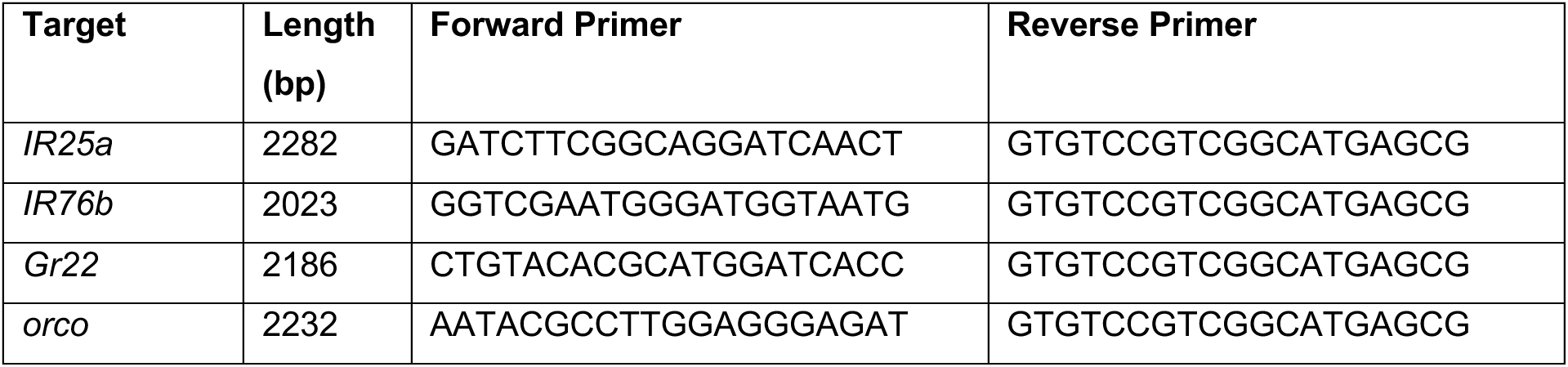
PCR primers used for verification of DRM cassette integration into each target *Anopheles gambiae* chemoreceptor gene.

## KEY RESOURCES TABLE

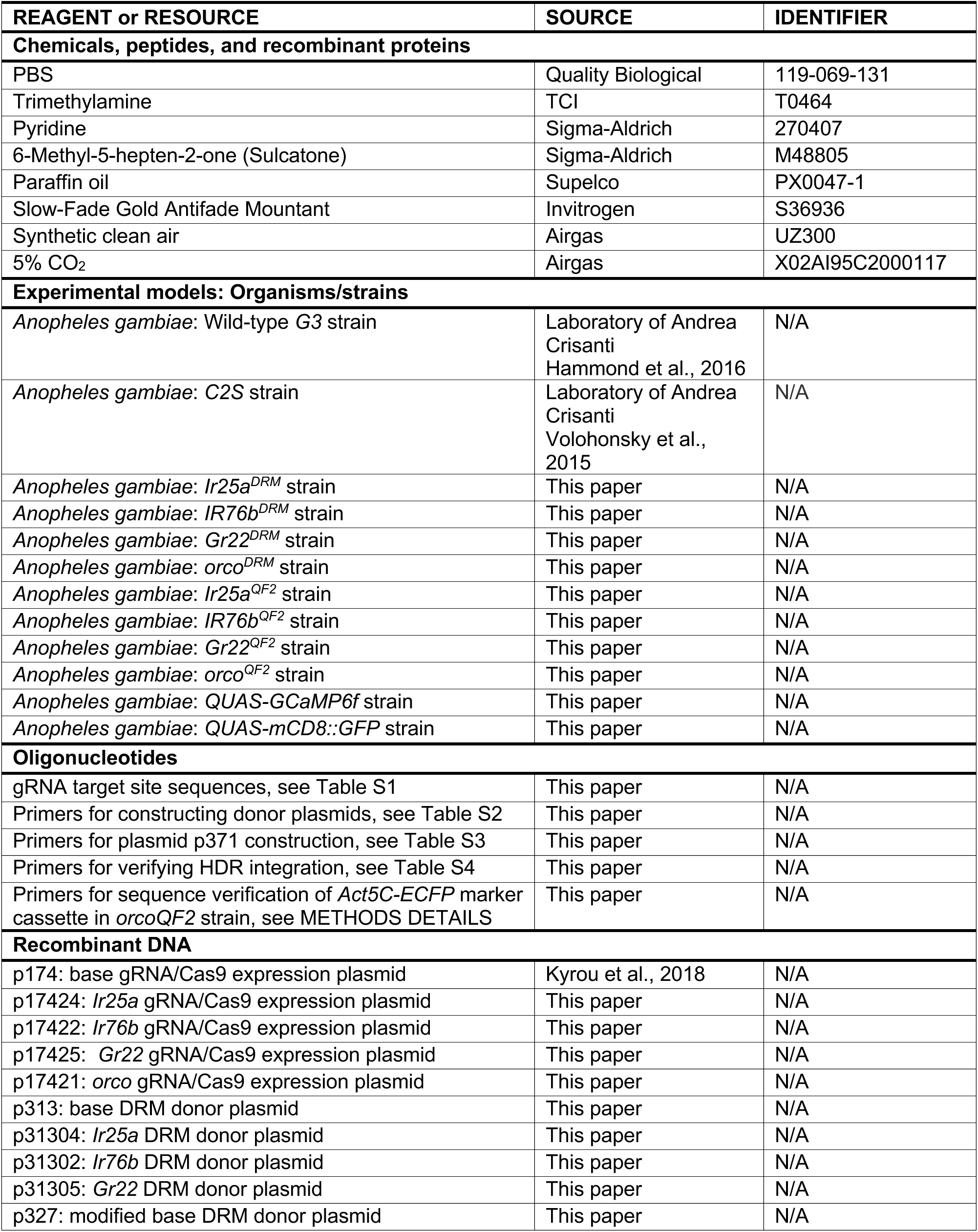
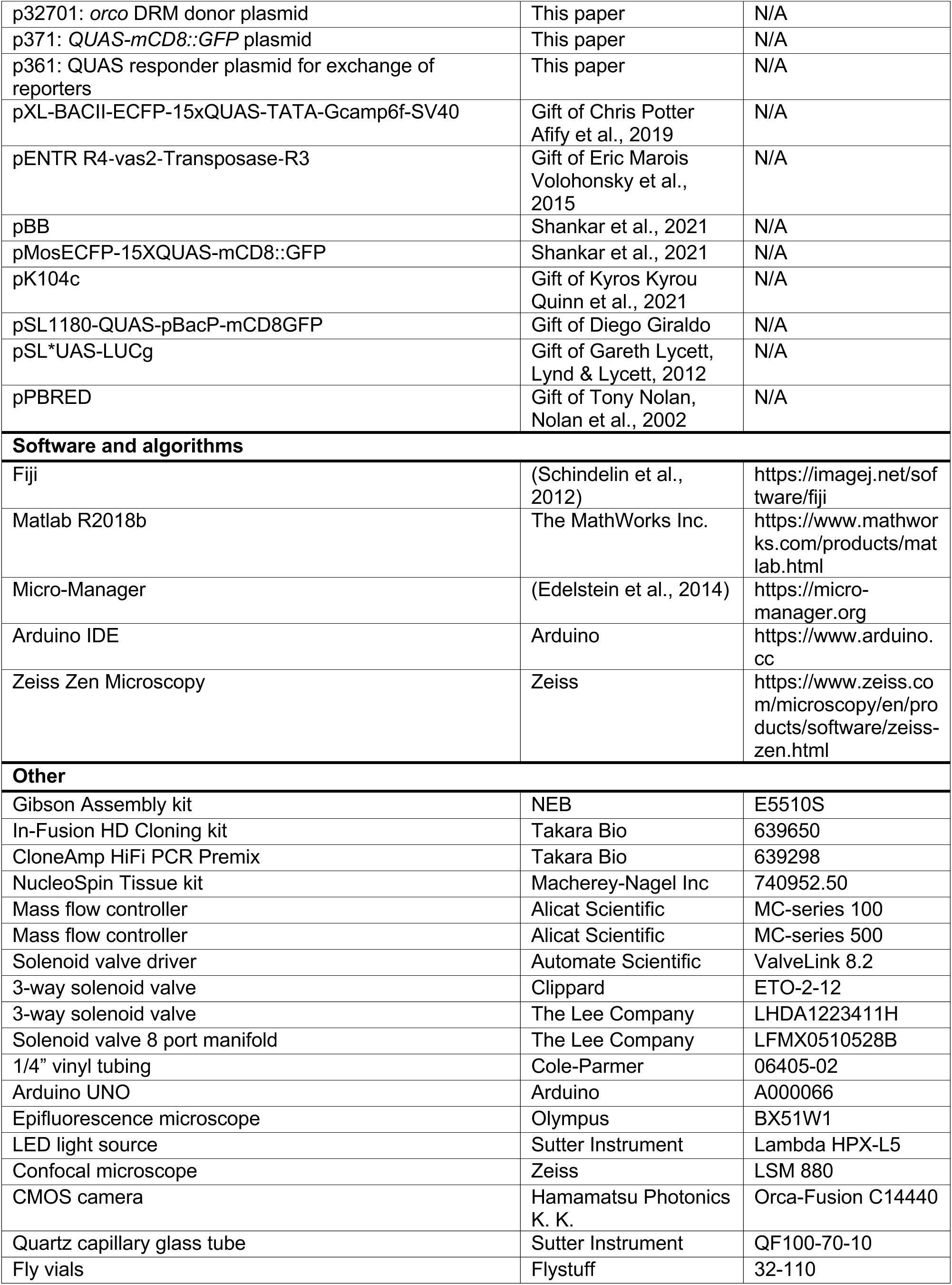
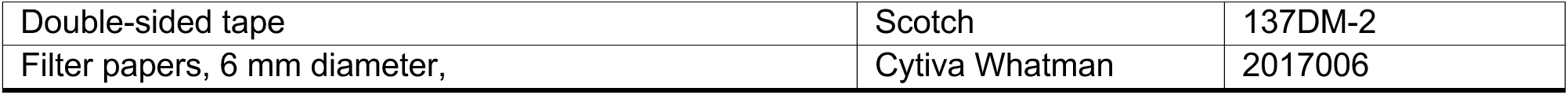

## Notes

### Competing Interest Statement

The authors have declared no competing interest.

## REFERENCES

Adolfi, A., Lynd, A., Lycett, G.J., and James, A.A. (2021). Site-Directed φC31-Mediated Integration and Cassette Exchange in *Anopheles* Vectors of Malaria. J Vis Exp. 168. doi: 10.3791/62146.

Afify, A., Betz, J.F., Riabinina, O., Lahondère, C., and Potter, C.J. (2019). Commonly Used Insect Repellents Hide Human Odors from Anopheles Mosquitoes. Curr. Biol. 29, 3669–3680.e5.

Athrey, G., Cosme, L. V., Popkin-Hall, Z., Pathikonda, S., Takken, W., and Slotman, M.A. (2017). Chemosensory gene expression in olfactory organs of the anthropophilic Anopheles coluzzii and zoophilic Anopheles quadriannulatus. BMC Genomics 18, 751.

Bateman, J.R., Lee, A.M., and Wu, C.T. (2006). Site-specific transformation of Drosophila via phiC31 integrase-mediated cassette exchange. Genetics. 173:769–777.

Benedict, M.Q., Hunt, C.M., Vella, M.G., Gonzalez, K.M., Dotson, E.M., and Matilda Collins, C. (2020). Pragmatic selection of larval mosquito diets for insectary rearing of Anopheles gambiae and Aedes aegypti. PLoS One 15, e0221838.

Berghammer, A.J., Klingler, M., and Wimmer, E.A. (1999). A universal marker for transgenic insects. Nature 402, 370–371.

Bernier, U.R., Booth, M.M., and Yost, R.A. (1999). Analysis of human skin emanations by gas chromatography/mass spectrometry. 1. Thermal desorption of attractants for the yellow fever mosquito (Aedes aegypti) from handled glass beads. Anal. Chem. 71, 1–7.

Bernier, U.R., Kline, D.L., Barnard, D.R., Schreck, C.E., and Yost, R.A. (2000). Analysis of human skin emanations by gas chromatography/mass spectrometry. 2. Identification of volatile compounds that are candidate attractants for the yellow fever mosquito (Aedes aegypti). Anal. Chem. 72, 747–756.

Besansky, N.J., Hill, C.A., and Costantini, C. (2004). No accounting for taste: Host preference in malaria vectors. Trends Parasitol. 20, 249–251.

Braks, M.A.H., Meijerink, J., and Takken, W. (2001). The response of the malaria mosquito, Anopheles gambiae, to two components of human sweat, ammonia and L-lactic acid, in an olfactometer. Physiol. Entomol. 26, 142–148.

Carey, A.F., Wang, G., Su, C.-Y., Zwiebel, L.J., and Carlson, J.R. (2010). Odorant reception in the malaria mosquito Anopheles gambiae. Nature 464, 66–71.

Cator, L.J., George, J., Blanford, S., Murdock, C.C., Baker, T.C., Read, A.F., and Thomas, M.B. (2013). “Manipulation” without the parasite: Altered feeding behaviour of mosquitoes is not dependent on infection with malaria parasites. Proc. R. Soc. B Biol. Sci. 280, 20130711.

Chen, T.W., Wardill, T.J., Sun, Y., Pulver, S.R., Renninger, S.L., Baohan, A., Schreiter, E.R., Kerr, R.A., Orger, M.B., Jayaraman, V., et al. (2013). Ultrasensitive fluorescent proteins for imaging neuronal activity. Nature 499, 295–300.

Constantini, C., Sagnon, N.F., della Torre, A., and Coluzzi, M. (1999). Mosquito Behavioral Aspects of Vector-Human Interactions in the Anopheles gambiae complex. Parasitologia 41, 209–217.

Cork, A., and Park, K.C. (1996). Identification of electrophysiologically-active compounds for the malaria mosquito, Anopheles gambiae, in human sweat extracts. Med. Vet. Entomol. 10, 269–276.

Coutinho-Abreu, I. V., Sharma, K., Cui, L., Yan, G., and Ray, A. (2019). Odorant ligands for the CO2 receptor in two Anopheles vectors of malaria. Sci. Rep. 9, 2549.

Dekker, T., Takken, W., and Braks, M.A.H. (2001). Innate preference for host-odor blends modulates degree of anthropophagy of Anopheles gambiae sensu lato (Diptera: Culicidae). J. Med. Entomol. 38, 868–871.

Edelstein, A.D., Tsuchida, M.A., Amodaj, N., Pinkard, H., Vale, R.D., and Stuurman, N. (2014). Advanced methods of microscope control using μManager software. J. Biol. Methods 1, 10.

Foster, W.A., and Takken, W. (2004). Nectar-related vs. human-related volatiles: behavioural response and choice by female and male Anopheles gambiae (Diptera: Culicidae) between emergence and first feeding. Bull. Entomol. Res. 94, 145–157.

Fox, A.N., Pitts, R.J., Robertson, H.M., Carlson, J.R., and Zwiebel, L.J. (2001). Candidate odorant receptors from the malaria vector mosquito Anopheles gambiae and evidence of down-regulation in response to blood feeding. Proc. Natl. Acad. Sci. 98, 14693–14697.

Garrett-Jones, C., Boreham, P.F.L., and Pant, C.P. (1980). Feeding habits of anophelines (Diptera: Culicidae) in 1971–78, with reference to the human blood index: A review. Bull. Entomol. Res. 70, 165–185.

Giraldo, D., Rankin-Turner, S., Corver, A., Tauxe, G.M., Gao, A.L., Jackson, D.M., Simubali, L., Book, C., Stevenson, J.C., Thuma, P.E., et al. (2023). Human scent guides mosquito thermotaxis and host selection under naturalistic conditions. Curr. Biol. 33, 1–16.

Hammond, A., Galizi, R., Kyrou, K., Simoni, A., Siniscalchi, C., Katsanos, D., Gribble, M., Baker, D., Marois, E., Russell, S., et. al. (2016). A CRISPR-Cas9 gene drive system targeting female reproduction in the malaria mosquito vector Anopheles gambiae. Nat Biotechnol 34: 78–83.

Han, K., Levine, M.S., and Manley, J.L. (1989). Synergistic activation and repression of transcription by Drosophila homeobox proteins. Cell. 56: 573–583

Herre, M., Goldman, O. V., Lu, T.-C., Caballero-Vidal, G., Qi, Y., Gilbert, Z.N., Gong, Z., Morita, T., Rahiel, S., Ghaninia, M., et al. (2022). Non-Canonical Odor Coding in the Mosquito. Cell 185, 3104–3123.

Hoermann, A., Tapanelli, S., Capriotti, P., Del Corsano, G., Masters, E.K.G., Habtewold, T., Christophides, G.K., and Windbichler, N. (2021). Converting endogenous genes of the malaria mosquito into simple non-autonomous gene drives for population replacement. Elife 10, e58791.

Huang, J., Miller, J.R., Chen, S.C., Vulule, J.M., and Walker, E.D. (2006). Anopheles gambiae (Diptera: Culicidae) oviposition in response to agarose media and cultured bacterial volatiles. J. Med. Entomol. 43, 498–504.

Knols, B.G.J., Van Loon, J.J.A., Cork, A., Robinson, R.D., Adam, W., Meijerink, J., De Jong, R., and Takken, W. (1997). Behavioural and electrophysiological responses of the female malaria mosquito Anopheles gambiae (Diptera: Culicidae) to Limburger cheese volatiles. Bull. Entomol. Res. 87, 151–159.

Konopka, J.K., Task, D., Poinapen, D., Potter, C.J. (2023) Neurogenetic identification of mosquito sensory neurons. iScience. 26: 106690.

Kyrou, K., Hammond, A.M., Galizi, R., Kranjc, N., Burt, A., Beaghton, A.K., Nolan, T., and Crisanti, A. (2018). A CRISPR-Cas9 gene drive targeting doublesex causes complete population suppression in caged Anopheles gambiae mosquitoes. Nat Biotechnol. 36:1062–1066.

Laursen, W.J., Budelli, G., Tang, R., Chang, E.C., Busby, R., Shankar, S., Gerber, R., Greppi, C., Albuquerque, R., and Garrity, P.A. (2023). Humidity sensors that alert mosquitoes to nearby hosts and egg-laying sites Article Humidity sensors that alert mosquitoes to nearby hosts and egg-laying sites. Neuron 1–14.

Liu, F., Ye, Z., Baker, A., Sun, H., and Zwiebel, L.J. (2020). Gene editing reveals obligate and modulatory components of the CO2 receptor complex in the malaria vector mosquito, Anopheles coluzzii. Insect Biochem. Mol. Biol. 127, 1–9.

Lu, T., Qiu, Y.T., Wang, G., Kwon, J.Y., Rutzler, M., Kwon, H.W., Pitts, R.J., van Loon, J.J.A., Takken, W., Carlson, J.R., et al. (2007). Odor Coding in the Maxillary Palp of the Malaria Vector Mosquito Anopheles gambiae. Curr. Biol. 17, 1533–1544.

Lynd, A., and Lycett, G.J. (2012). Development of the bi-partite Gal4-UAS system in the African malaria mosquito, *Anopheles gambiae*. PLoS One. 7: e31552.

Matthews BJ, Dudchenko O, Kingan SB, Koren S, Antoshechkin I, Crawford JE, Glassford WJ, Herre M, Redmond SN, Rose NH et al. (2018) Improved reference genome of Aedes aegypti informs arbovirus vector control. Nature. 563: 501–507.

Markstein, M., Pitsouli, C., Villalta, C., Celniker, S.E., and Perrimon, N. (2008). Exploiting position effects and the gypsy retrovirus insulator to engineer precisely expressed transgenes. Nat. Genet. 40, 476–483.

Martínez-Lozano, P. (2009). Mass spectrometric study of cutaneous volatiles by secondary electrospray ionization. Int. J. Mass Spectrom. 282, 128–132.

McIver, S.B. (1982). Sensilla mosquitoes (Diptera: Culicidae). J. Med. Entomol. 19, 489–535.

Meijerink, J., and Van Loon, J.J.A. (1999). Sensitivities of antennal olfactory neurons of the malaria mosquito, Anopheles gambiae, to carboxylic acids. J. Insect Physiol. 45, 365–373.

Mozūraitis, R., Hajkazemian, M., Zawada, J.W., Szymczak, J., Pålsson, K., Sekar, V., Biryukova, I., Friedländer, M.R., Koekemoer, L.L., Baird, J.K., et al. (2020). Male swarming aggregation pheromones increase female attraction and mating success among multiple African malaria vector mosquito species. Nat. Ecol. Evol. 4, 1395–1401.

Mukabana, W.R., Takken, W., Coe, R., and Knols, B.G.J. (2002). Host-specific cues cause differential attractiveness of Kenyan men to the African malaria vector Anopheles gambiae. Malar. J. 1, 17.

Ni, L. (2021). The Structure and Function of Ionotropic Receptors in Drosophila. Front. Mol. Neurosci. 13, 1–11.

Nikbakhtzadeh, M.R., Terbot II, J.W., Otienoburu, P.E., and Foster, W.A. (2014). Olfactory basis of floral preference of the malaria vector Anopheles gambiae (Diptera: Culicidae) among common African plants. J. Vector Ecol. 39, 372–383.

Nolan, T., Bower, T.M., Brown, A.E., Crisanti, A., Catteruccia, F. (2002) *piggyBac*-mediated germline transformation of the malaria mosquito *Anopheles stephensi* using the red fluorescent protein dsRED as a selectable marker. J. Biol. Chem. 277: 8759–62.

Nyasembe, V.O., Teal, P.E., Mukabana, W.R., Tumlinson, J.H., and Torto, B. (2012). Behavioural response of the malaria vector Anopheles gambiae to host plant volatiles and synthetic blends. Parasites and Vectors 5, 234.

O’Brochta, D.A., Pilitt, K.L., Harrell, R.A., Aluvihare, C., and Alford, R.T. (2012). Gal4-based Enhancer-Trapping in the Malaria Mosquito Anopheles stephensi. G3 2, 1305–1315.

Omondi, A.B., Ghaninia, M., Dawit, M., Svensson, T., and Ignell, R. (2019). Age-dependent regulation of host seeking in Anopheles coluzzii. Sci. Rep. 9, 1–9.

Omondi, B.A., Majeed, S., and Ignell, R. (2015). Functional development of carbon dioxide detection in the maxillary palp of Anopheles gambiae. J. Exp. Biol. 218, 2482–2488.

Pfeiffer, B.D., Ngo, T.T.B., Hibbard, K.L., Murphy, C., Jenett, A., Truman, J.W., and Rubin, G.M. (2010). Refinement of tools for targeted gene expression in Drosophila. Genetics 186, 735– 755.

Pfeiffer, B.D., Truman, J.W., and Rubin, G.M. (2012). Using translational enhancers to increase transgene expression in Drosophila. PNAS 109, 6626–6631.

Pitts, R.J., Rinker, D.C., Jones, P.L., Rokas, A., and Zwiebel, L.J. (2011). Transcriptome profiling of chemosensory appendages in the malaria vector Anopheles gambiae reveals tissue- and sex-specific signatures of odor coding. BMC Genomics 12, 271.

Pitts, R.J., Derryberry, S.L., Zhang, Z., and Zwiebel, L.J. (2017). Variant Ionotropic Receptors in the Malaria Vector Mosquito Anopheles gambiae Tuned to Amines and Carboxylic Acids. Sci. Rep. 7, 40297.

Qiu, Y.T., and van Loon, J.A. (2010). Olfactory physiology of blood-feeding vector mosquitoes. In Olfaction in Vector-Host Interactions, pp. 39–61.

Qiu, Y.T., Smallegange, R.C., Hoppe, S., van Loon, J.J.A., Bakker, E.-J., and Takken, W. (2004). Behavioural and electrophysiological responses of the malaria mosquito Anopheles gambiae Giles sensu stricto (Diptera: Culicidae) to human skin emanations. Med. Vet. Entomol. 18, 429–438.

Qiu, Y.T., van Loon, J.J.A., Takken, W., Meijerink, J., and Smid, H.M. (2006). Olfactory coding in antennal neurons of the malaria mosquito, Anopheles gambiae. Chem. Senses 31, 845–863.

Quinn, C., Anthousi, A., Wondji, C., and Nolan, T. (2021). CRISPR-mediated knock-in of transgenes into the malaria vector *Anopheles funestus*. G3 (Bethesda). 11: jkab201. doi: 10.1093/g3journal/jkab201.

Raji, J.I., Konopka, J.K., and Potter, C.J. (2023). A spatial map of antennal-expressed ionotropic receptors in the malaria mosquito. Cell Rep. 42, 112101.

Rankin-Turner, S., and McMeniman, C.J. (2022). A headspace collection chamber for whole body volatilomics. Analyst 147, 5210.

Riabinina, O., Task, D., Marr, E., Lin, C.-C., Alford, R., O’Brochta, D.A., and Potter, C.J. (2016). Organization of olfactory centres in the malaria mosquito Anopheles gambiae. Nat. Commun. 7, 13010.

Rinker, D.C., Pitts, R.J., Zhou, X., Suh, E., Rokas, A., and Zwiebel, L.J. (2013). Blood meal-induced changes to antennal transcriptome profiles reveal shifts in odor sensitivities in Anopheles gambiae. Proc. Natl. Acad. Sci. 110, 8260–8265.

Saveer, A.M., Pitts, R.J., Ferguson, S.T., and Zwiebel, L.J. (2018). Characterization of Chemosensory Responses on the Labellum of the Malaria Vector Mosquito, Anopheles coluzzii. Sci. Rep. 8, 5656.

Schindelin, J., Arganda-Carreras, I., Frise, E., Kaynig, V., Longair, M., Pietzsch, T., Preibisch, S., Rueden, C., Saalfeld, S., Schmid, B., et al. (2012). Fiji: An open-source platform for biological-image analysis. Nat. Methods 9, 676–682.

Schinko JB, Weber M, Viktorinova I, Kiupakis A, Averof M, Klingler M, Wimmer EA, Bucher G. (2010) Functionality of the GAL4/UAS system in Tribolium requires the use of endogenous core promoters. BMC Dev Biol. 19;10:53.

Scott, T.W., and Takken, W. (2012). Feeding strategies of anthropophilic mosquitoes result in increased risk of pathogen transmission. Trends Parasitol. 28, 114–121.

Shankar, S., Tauxe, G.M., Spikol, E.D., Li, M., Akbari, O.S., Giraldo, D., and McMeniman, C.J. (2021). Synergistic coding of carbon dioxide and a human sweat odorant in the mosquito brain. bioRxiv. doi: https://doi.org/10.1101/2020.11.02.365916365916v2

Smallegange, R.C., Knols, B.G.J., and Takken, W. (2010). Effectiveness of Synthetic Versus Natural Human Volatiles as Attractants for *Anopheles gambiae* (Diptera: Culicidae) Sensu Stricto. J. Med. Entomol. 47, 338–344.

Spitzen, J., Spoor, C.W., Grieco, F., ter Braak, C., Beeuwkes, J., van Brugge, S.P., Kranenbarg, S., Noldus, L.P.J.J., van Leeuwen, J.L., and Takken, W. (2013). A 3D Analysis of Flight Behavior of Anopheles gambiae sensu stricto Malaria Mosquitoes in Response to Human Odor and Heat. PLoS One 8, e62995.

Stern, D.L. (2022). Transgenic tools for targeted chromosome rearrangements allow construction of balancer chromosomes in non-melanogaster Drosophila species. G3 Genes, Genomes, Genet. 12, jkac030.

Sun, H., Liu, F., Ye, Z., Baker, A., and Zwiebel, L.J. (2020). Mutagenesis of the orco odorant receptor co-receptor impairs olfactory function in the malaria vector Anopheles coluzzii. Insect Biochem. Mol. Biol. 127, 103497.

Takken, W., and Knols, B.G.J. (1999). Odor-Mediated Behavior of Afrotropical Malaria Mosquitoes. Annu. Rev. Entomol. 44, 131–157.

Takken, W., and Verhulst, N.O. (2013). Host preferences of blood-feeding mosquitoes. Annu. Rev. Entomol. 58, 433–453.

Takken, W., Van Loon, J.J.A., and Adam, W. (2001). Inhibition of host-seeking response and olfactory responsiveness in Anopheles gambiae following blood feeding. J. Insect Physiol. 47, 303–310.

Task, D., Lin, C.-C., Vulpe, A., Afify, A., Ballou, S., Brbic, M., Schlegel, P., Raji, J., Jefferis, G., Li, H., et al. (2022). Chemoreceptor co-expression in Drosophila melanogaster olfactory neurons. Elife 11, 1–69.

Volohonsky, G., Terenzi, O., Soichot, J., Naujoks, D.A., Nolan, T., Windbichler, N., Kapps, D., Smidler, A.L., Vittu, A., Costa, G., et al. (2015). Tools for Anopheles gambiae transgenesis. G3 Genes, Genomes, Genet. 5, 1151–1163.

Wang, G., Carey, A.F., Carlson, J.R., and Zwiebel, L.J. (2010). Molecular basis of odor coding in the malaria vector mosquito Anopheles gambiae. Proc. Natl. Acad. Sci. U. S. A. 107, 4418– 4423.

Ye, Z., Liu, F., Sun, H., Baker, A., and Zwiebel, L.J. (2022). Discrete Roles of the Ir76b Ionotropic Co-Receptor Impact Olfaction, Blood Feeding, and Mating in the Malaria Vector Mosquito Anopheles coluzzii. PNAS 119, e2112385119.

Zhao, Z., Tian, D., and McBride, C.S. (2021). Development of a pan-neuronal genetic driver in Aedes aegypti mosquitoes. Cell Reports Methods 1, 100042.

Zhao, Z., Zung, J.L., Hinze, A., Kriete, A.L., Iqbal, A., Younger, M.A., Matthews, B.J., Merhof, D., Thiberge, S., Ignell, R., et al. (2022). Mosquito brains encode unique features of human odour to drive host seeking. Nature 605, 706–712.

